# Continental-scale computer vision models reveal generalizable patterns and pitfalls for urban tree inventories with street-view images

**DOI:** 10.1101/2025.10.09.681424

**Authors:** Thomas A. Lake, Brit B. Laginhas, Brennen T. Farrell, Ross K. Meentemeyer, Chris M. Jones

**Author notes:** Corresponding authors at: North Carolina State University, Center for Geospatial Analytics, 5112 Jordan Hall, 2800 Faucette Dr, Raleigh, NC 27695, *Email addresses. Co-first authors (Thomas A. Lake, Brit B. Laginhas).

## Abstract

Accurate, up-to-date catalogs of urban tree populations are crucial for quantifying ecosystem services and enhancing the quality of life in cities. However, mapping tree species cost-effectively remains challenging. In response, remote sensing researchers are developing general-purpose tools to survey plant populations across broad spatial scales. In this study, we developed computer vision models to detect, classify, and map 100 tree genera across 23 cities in North America using Google Street View (GSV) and iNaturalist images. We validated our predictions in independent portions of each city. We then compared our predictions to existing street tree records to evaluate the spatial context of errors using generalized linear mixed-effects models. Our computer vision models identified most ground-truthed street trees (67.1%). Performance varied across the 23 cities (67.4% ± 9.3%) and 100 genera (50.9% ± 23.0%) and improved denser street-view coverage, simpler stand structure, and greater training representation, particularly from the focal city. We found that genus classification performed better in continental cities with lower relative diversity, and that seasonal changes in the appearance of trees provided visual cues that moderate classification rates. Using widely available street-level imagery is a generalizable and promising avenue for mapping tree distributions across urban environments.

## 1. Introduction

Urban forests, comprising all trees in urban settings, influence the livability and quality of life in cities (Edgar et al., 2021; Nowak and Dwyer, 2007; Roy et al., 2012). These forests provide necessary ecosystem services, including climate regulation, stormwater management, and pollution sequestration, valued at over $18 billion annually in the United States (Nowak and Dwyer, 2007; Nowak and Greenfield, 2018). Understanding tree distribution, species composition, and health is essential for quantifying and managing these benefits (Ma et al., 2021; McPherson et al., 2016). Traditionally, field surveys are used to document forest conditions, but collecting data on individual trees is labor-intensive and quickly becomes outdated due to urban development and pest or disease impacts (Nielsen et al., 2014; Pretzsch et al., 2017; Raum et al., 2023; Tubby and Webber, 2010). As such, there is a growing need for efficient, scalable approaches to monitor urban forests.

Tree inventories remain the primary means by which cities document the location and status of individual trees (Nielsen et al., 2014; Nowak et al., 2008; Roman et al., 2013). Inventories are typically conducted by professional arborists, and their accuracy is crucial for quantifying ecosystem services and forecasting pest or pathogen risk (Hargrave et al., 2022; Jones et al., 2021; Jones et al., 2022; Ma et al., 2021; Miller et al., 2015). However, few communities maintain updated inventories (Hargrave et al., 2022). National programs such as the USDA Urban Forest Inventory and Analysis provide systematic plot-based estimates of urban forest characteristics (Edgar et al., 2021), but not a full census. Consequently, many cities rely on opportunistic surveys and community science observations to supplement records and track forest conditions (Bloniarz, 1996; Crown et al., 2018; Roman et al., 2017). Researchers are increasingly turning to remote sensing and machine learning for more comprehensive and timely assessments of urban trees.

Advances in remote sensing have improved our ability to map and monitor urban forests (Fassnacht et al., 2016; Ward and Johnson, 2007). Tree species with distinct morphological, phenological, and spectral traits can be identified from aerial and satellite imagery (Fassnacht et al., 2016; Nagendra, 2001). Numerous studies have combined high-resolution images with light detection and ranging (LiDAR) to inventory urban tree species (Hartling et al., 2019; Pu et al., 2018; Pu and Landry, 2020; Zhang and Hu, 2012). Yet, extending these approaches beyond individual cities remains limited by the cost of fine-resolution data and inconsistent image quality, acquisition timing, and coverage (Velasquez-Camacho et al., 2021). As a result, these methods are typically applied at local scales (e.g., individual parks, neighborhoods, or cities) and rarely extended across multiple urban areas for broader comparison (Beery et al., 2022; Li et al., 2019; Velasquez-Camacho et al., 2021).

Street-level imagery provides a complementary perspective by offering ground-level views of urban trees (Berland and Lange, 2017; Berland et al., 2019; Roberts et al., 2019; Seiferling et al., 2017). Google Street View (GSV) captures high-resolution panoramas every 10-20 meters along roads worldwide (Anguelov et al., 2010; Biljecki and Ito, 2021). Similar platforms (e.g., Mapillary; https://www.mapillary.com/), provide analogous views with crowdsourced imagery. These images capture canopy and tree characteristics, including leaf arrangement, branching, and bark texture, that are typically obscured in aerial imagery. Yet, most studies examine individual neighborhoods or cities (Branson et al., 2018; Capecchi et al., 2023; Choi et al., 2022; Liu et al., 2023; Lumnitz et al., 2021; Velasquez-Camacho et al., 2024; Wegner et al., 2016), and it remains unclear how well street-level surveys can generalize across broader regions.

Machine learning models are increasingly used to detect and classify plant species from remotely-sensed imagery, including aerial, satellite, and street-level sources (Hartling et al., 2019; Lake et al., 2022; Lumnitz et al., 2021; Ventura et al., 2024). These models can achieve high accuracy when trained and tested within the same region but often perform less reliably when applied to new areas (Beery et al., 2022; Marconi et al., 2022). Model generalizability is constrained by environmental and seasonal variation, and by factors related to image and site characteristics (Capecchi et al., 2023; Choi et al., 2022; Kim and Jang, 2023; Liu et al., 2023; Ventura et al., 2024; Velasquez-Camacho et al., 2024). As a result, developing models that generalize reliably across regions remains a challenge (Arevalo-Ramirez et al., 2024; Beery et al., 2022; Van Horn and Perona, 2017).

In this study, we developed computer vision models to detect, classify, and map urban street trees across 23 cities in the United States and Canada using GSV and iNaturalist images. We trained tree detection and classification models and evaluated their performance using independent test splits that captured both cross-city and within-city generalization. Our objectives were to (1) examine how variation in urban tree diversity across cities influences image classification accuracy, (2) test whether geography and the month of GSV image capture explain variation in accuracy, and (3) determine how ecological conditions and street-view image coverage influence model errors and generalization across cities. We used model predictions to generate city-level tree inventories and fitted generalized linear mixed-effect models to evaluate when and where predictions aligned with known street trees.

## 2. Materials and methods

### 2.1 Study cities

We examined public street trees in 23 U.S. and Canadian cities. These cities have existing tree inventories, GSV imagery, and span diverse geographic, ecological, and socioeconomic contexts. Cities with smaller populations in northern, continental climates (e.g., Sioux Falls, South Dakota, and Kitchener, Ontario) have lower tree diversity, mostly consisting of native deciduous and coniferous species (Table 1). In contrast, metropolitan cities, such as Los Angeles, California, and New York City, New York, exhibit higher tree diversity with a greater proportion of non-native and ornamental species (Table 1) (Love et al., 2022; Ma et al., 2020; McCoy et al., 2022). Variation in urban populations, environmental conditions, and management practices shapes the species composition and diversity of urban forests (Roman et al., 2018), presenting opportunities and challenges for tree inventories with street-level imagery.

**Table 1.** Characteristics of the 23 study cities. For each city, we report latitude and longitude coordinates; its assigned geographic region (determined via principal component analysis of WorldClim v2.1 bioclimatic variables); tree diversity (genus richness and the Shannon diversity index); human population (2020 U.S. Census or the 2021 Canadian Census); mean annual temperature (in °C; BIO1); and total annual precipitation (in mm; BIO12).

| City | Location | Geographic Region | Genus Richness | Shannon Diversity Index | Human Population size | Mean Annual Temp (°C) | Annual Precipitation (mm) |
| --- | --- | --- | --- | --- | --- | --- | --- |
| Bloomington, IN | 39.165, -86.536 | East | 47 | 2.99 | 79,077 | 11.6 | 90.57 |
| Boulder, CO | 40.025, -105.277 | Central | 49 | 2.95 | 108,123 | 9.7 | 79.86 |
| Buffalo, NY | 42.885, -78.874 | East | 57 | 2.74 | 277,551 | 8.4 | 91.19 |
| Calgary, AB | 51.044, -114.073 | Central | 28 | 2.4 | 1,481,806 | 3.4 | 90.23 |
| Cambridge, ON | 43.405, -80.329 | East | 51 | 2.66 | 138,479 | 7.0 | 94.46 |
| Charlottesville, VA | 38.032, -78.482 | East | 48 | 3.07 | 46,469 | 13.5 | 81.32 |
| Columbus, OH | 39.963, -83.002 | East | 64 | 3.26 | 906,418 | 10.9 | 90.9 |
| Cupertino, CA | 37.317, -122.046 | Southwest | 71 | 3.18 | 60,111 | 14.8 | 37.94 |
| Denver, CO | 39.737, -104.991 | Central | 64 | 3.4 | 717,606 | 10.2 | 83.5 |
| Edmonton, AB | 53.536, -113.495 | Central | 28 | 2.42 | 1,418,118 | 2.8 | 108.01 |
| Kitchener, ON | 43.447, -80.481 | East | 26 | 2.11 | 256,885 | 6.6 | 94.42 |
| Los Angeles, CA | 34.052, -118.253 | Southwest | 82 | 4.16 | 3,895,848 | 18.3 | 33.99 |
| Montreal, QC | 45.512, -73.586 | East | 56 | 3.14 | 1,762,949 | 6.5 | 108.04 |
| New York, NY | 40.720, -74.001 | East | 61 | 3.70 | 8,740,292 | 12.0 | 87.57 |
| Pittsburgh, PA | 40.440, -79.995 | East | 64 | 2.92 | 302,779 | 10.6 | 87.58 |
| San Francisco, CA | 37.768, -122.41 | Southwest | 83 | 3.99 | 870,518 | 14.1 | 22.48 |
| San Jose, CA | 37.331, -121.88 | Southwest | 90 | 4.02 | 1,009,319 | 15.4 | 40.72 |
| Santa Monica, CA | 34.023, -118.48 | Southwest | 60 | 3.54 | 92,913 | 17.2 | 26.28 |
| Seattle, WA | 47.604, -122.333 | Northwest | 76 | 3.70 | 740,565 | 10.9 | 48.39 |
| Sioux Falls, SD | 43.542, -96.731 | Central | 35 | 1.78 | 193,437 | 7.4 | 113.5 |
| Surrey, BC | 49.187, -122.846 | Northwest | 59 | 3.24 | 568,322 | 9.5 | 50.36 |
| Vancouver, BC | 49.279, -123.123 | Northwest | 66 | 3.35 | 662,248 | 9.9 | 52.91 |
| Washington, DC | 38.898, -77.038 | East | 56 | 3.38 | 689,545 | 13.2 | 85.44 |

### 2.2 Data collection and image filtering

#### 2.2.1 Selecting focal genera

We selected 100 genera to detect, classify, and geolocate across 23 cities (Table S1). We worked at the genus level to avoid distinguishing between numerous varieties, hybrids, and subspecies commonly found in urban areas. Tree species and genus diversity follows a long-tailed distribution, where a few taxa are abundant and most are infrequent or rare (Love et al., 2022; Ma et al., 2020; McCoy et al., 2022). Our selected genera account for over 95% of all genera reported in the 23 inventories. These include taxa that are widely distributed (e.g., *Acer* [maple], *Fraxinus* [ash]), regionally abundant (e.g., *Magnolia* [magnolia], *Picea* [spruce]), and occasionally found as non-native or ornamental species (e.g., *Ailanthus* [tree of heaven], *Lagerstroemia* [crape myrtle]).

#### 2.2.2 AutoArborist tree genera images

We used a comprehensive GSV dataset to develop computer vision models that detect and classify tree genera (Figure 1). Beery et al. (2022) created the AutoArborist dataset (v0.16) by co-locating GSV images with tree inventory records across 23 North American cities. This dataset includes approximately 1.2 million GSV images of over 300 genera. The images (dimensions: 768 x 1152 pixels) also contain metadata, including capture location (latitude and longitude), capture date (month and year), and annotated bounding boxes of individual trees.

**Figure 1.**
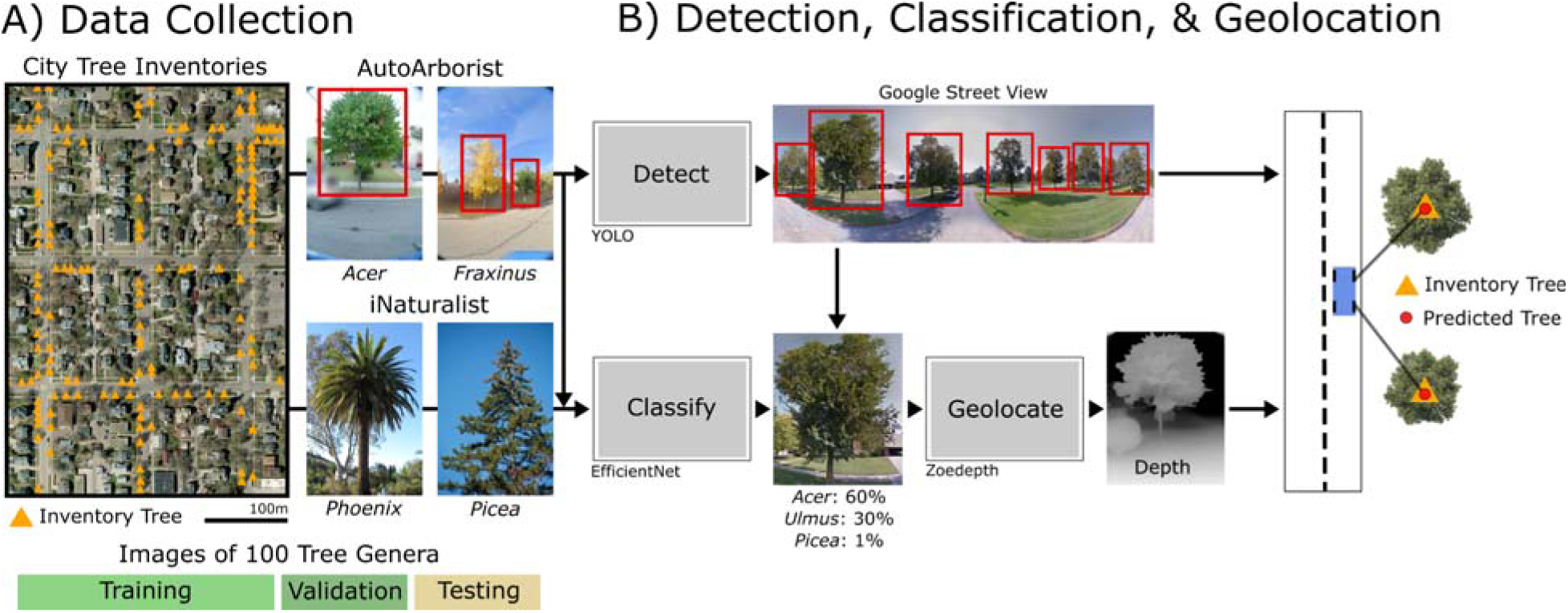
Computer vision-based tree inventories. A) Data collection: Images for 100 tree genera were sourced from AutoArborist (Google Street View [GSV] images from 23 cities in United States and Canada) and research-grade observations on iNaturalist.org. B) Detection, classification, and geolocation: An object detection (YOLOv5x) model and image classification (EfficientNetV2S) model identify tree genera in GSV images. Next, a depth estimation model (ZoeDepth) assigns geographic coordinates to predicted trees. The workflow is evaluated on spatially independent holdout areas within each city to assess out-of-sample generalizability.

#### 2.2.3 iNaturalist tree genera images

We augmented AutoArborist with iNaturalist images, a platform hosting millions of user-annotated plant images. We downloaded all available ‘research-grade’ images of our 100 genera captured in North America from their webserver (https://registry.opendata.aws/inaturalist-open-data; accessed August 1st, 2024), yielding approximately 3.6 million images.

#### 2.2.4 Filtering iNaturalist images

iNaturalist records are community-sourced with images taken from a wide range of viewpoints. Many are close-range (i.e., within one meter) photos of leaves, twigs, flowers, and fruits. While these features are important for taxonomic identification, they do not match the street-level perspective of GSV, where trunk and canopy are visible. Prior work shows that mixing close-ups with broader views (e.g., UAV or satellite) introduces visual inconsistencies that confuse computer vision models and degrade performance (Beery et al., 2022; Soltani et al., 2022; Soltani et al., 2024). Accordingly, we prioritized iNaturalist images with perspectives more consistent with GSV.

We applied a Contrastive Language-Image Pre-Training (CLIP) deep learning model to filter iNaturalist images (Radford et al., 2021). This model was trained on millions image-text pairs and ranks images based on alignment to a text prompt (Radford et al., 2021). We queried “a photo of a large mature tree in an urban landscape” to retain images consistent with the GSV perspective (Figure S1). We capped the number of retained images per genus to limit class imbalance. For genera with more than 10,000 images (e.g., *Acer* [maple]; 303,000 images), we kept the top 1,000 images based on prompt similarity. For genera with fewer than 10,000 images, we kept the top 10% of images. This ensured iNaturalist served as a supplement rather than a replacement for AutoArborist imagery. The curated subset totaled 59,132 images, which we incorporated into the training dataset for the tree classification model (see Section 2.3.2; Figure S1; Table S1).

### 2.3 Tree detection, classification, and geolocation models

#### 2.3.1 Tree detection

We developed an object detection model (YOLO; You Only Look Once) to identify trees in GSV images (Figure 1). We selected the YOLOv5 model (Jocher et al., 2022; Redmon et al., 2016) over R-CNN (Girshick et al., 2014) and Faster R-CNN (Ren et al., 2017) for its efficiency, accuracy, and ability to detect multi-scale objects (Terven and Cordova-Esparza, 2023). Previous studies likewise used YOLO variants (e.g., YOLOv3) to detect trees in street-level images (Choi et al., 2022; Liu et al., 2023).

We trained the object detection model on bounding box annotated tree images from AutoArborist (Beery et al., 2022). Because our goal was to build a single-class (tree) object detector that generalizes across all cities, we randomly sampled images across all genera to capture variation in tree appearance and image conditions. We sampled 750,000 images for training, 75,000 images for validation to fine-tune model parameters during training, and 75,000 images for testing. To improve generalization, we applied image augmentations (random horizontal flips, rotations, color shifts) during training to vary the appearance, scale, and illumination.

We trained for 25 epochs and evaluated performance on the test dataset using precision and recall, calculated at an Intersection over Union (IoU) threshold of 0.5. IoU measures the overlap between predicted and annotated boxes. Detections with an IoU of at least 0.5 were counted as true positives; otherwise they were classified as false positives. We selected this threshold to account for annotation inaccuracies in the AutoArborist dataset. For example, defining bounding boxes for complex objects such as trees is subjective (Everingham et al., 2010).

We calculated precision and recall to evaluate model performance. Precision quantifies resistance to false positives and is calculated as the number of true positives divided by the total predicted detections. Recall measures the ability to correctly detect trees, and is calculated as true positives divided the total number of annotated trees. Further details on the model’s architecture, training, and evaluation are provided in Supplementary Methods 1.1.

#### 2.3.2 Tree classification

We developed a convolutional neural network (CNN) to classify tree genera in GSV images (Figure 1). CNNs deliver state-of-the-art performance in image classification tasks across disciplines (Kattenborn et al., 2021; Krizhevsky et al., 2012). We selected EfficientNetV2-S for its balance of network complexity and computational efficiency (Tan and Le, 2021).

We trained and evaluated the image classification model using the 100 genera sourced from AutoArborist and iNaturalist (range: 793 images for *Larix* [larch] to 45,743 for *Acer* [maple]; Table S1). To assess generalization within and across cities, we applied a two-stage splitting strategy. For each city, we split the dataset into two spatially distinct subsets (Figure S2). Approximately 70% of images (N = 789,820) were allocated to model development, and 30% (N = 356,341) were withheld as a geographically independent test set for final evaluation. This spatial partitioning prevented geographic overlap between training and testing images.

Within the 70% development set, we further partitioned images into model training (80%), validation (10%), and internal test (10%) sets using random sampling. The validation set was used to monitor performance and tune model parameters during training, while the internal test set served as a holdout to assess cross-city classification performance (see Section *2.4.1*).

We trained the classification model for up to 100 epochs and terminated training when validation loss did not decrease after 10 consecutive epochs to limit overfitting. To improve generalization to unseen images, we applied data augmentations (random horizontal flips, rotations, and color shifts) during training. Model performance was assessed after training by computing recall (true positive rate), precision (proportion of predicted positives that are correct), and F1 score (a performance metric that accounts for both model recall and precision) for each genus on the 30% geographically stratified test split. Additional details on the model architecture, training, and evaluation are provided in Supplementary Methods 1.2.

#### 2.3.3. Tree classification model interpretation

Image classification performance generally improves with additional training data (Goodfellow et al., 2016; Hestness et al., 2017). To examine this effect across genera, we calculated the Spearman rank correlation coefficient between the number of training images per genus and test performance (the F1 score calculated on the 30% geographically stratified test split).

To understand which visual features underlie correct classifications, we applied gradient-weighted class activation maps (Grad-CAMs) (Selvaraju et al., 2017). Class activation mapping highlights image regions most influential to a model’s prediction (Zhou et al. 2016). Previous work using Grad-CAM indicates that bark and trunk features are important for species classification (Kim et al., 2022). We calculated Grad-CAMs from the penultimate convolutional layer in the image classification model for a random sample of 100 images per genus drawn from the AutoArborist 30% geographic testing split. We then visually inspected the Grad-CAMs to confirm that the model emphasized biologically meaningful features, such as leaves, branches, fruit, and crown structure, during inference.

#### 2.3.4. Tree geolocation

We geolocated each detected and classified tree in the GSV image using the image’s latitude and longitude, the angle of the tree, and the estimated camera-to-tree distance. We estimated distances with ZoeDepth, a monocular depth estimation model (Bhat et al. 2023). Monocular depth estimation infers object distances from single-color images (Godard et al., 2018) and have been used to approximate distances between GSV images and urban objects, including trees (Krylov et al., 2018; Liu et al., 2023; Lumnitz et al., 2021). We employed the pre-trained ZoeD-NK model, which was optimized for outdoor street-level imagery (Bhat et al., 2023). For each tree, the camera-to-tree distance was calculated as the median predicted depth within its bounding box. To mitigate distortions from the GSV panoramic projection, we multiplied each depth estimate by three, following Lumnitz et al. (2021).

### 2.4 Impact of diversity, image timing, and geography on genus classification

#### 2.4.1 Effect of diversity

We examined how inter-city variation in urban tree diversity influences image classification rates. Within each city, genus diversity follows a long-tailed distribution in which a few genera are common and many are rare (Love et al. 2022; Ma et al. 2020). This class imbalance biases model predictions, typically yielding higher accuracy for common genera than for rare ones (Beery et al., 2022; Van Horn and Perona, 2017).

To characterize urban tree diversity, we calculated three metrics for each city from the full AutoArborist Inventory (Table 1): richness (number of genera), evenness (0-1; relative abundances of genera), and the Shannon diversity index (integrates richness and evenness; Hill, 1973). High richness with low evenness indicates many genera with a few dominant taxa, whereas high richness with high evenness reflects a balanced genus composition. Shannon values typically range from 1.5 to 4.0, with higher values indicating greater richness and a more even genus distribution (Love et al., 2022). All metrics were computed in R using the vegan package (version 2.6-4; Oksanen et al., 2013).

To evaluate inter-city model performance, we used the internal 10% test set (N = 74,631 images) defined during the initial model development split (*Section 2.3.2*). This test set provides representative samples of each city with respect to richness, evenness and diversity. We then calculated the weighted F1 score across genera for each city. This metric is the harmonic mean of precision and recall, averaged across genera and weighted by the number of samples in each genus. Weighting accounts for class imbalance in each city by giving greater weight to common genera while retaining contributions to rare ones.

To test whether classification performance was associated with urban tree diversity, we fitted separate beta regression models for each diversity metric using the R package ‘betareg’ (Cribari-Neto and Zeileis, 2010). Model diagnostics confirmed that assumptions were met. One city, Kitchener, Ontario, was excluded from all regressions as an outlier due to anomalously low classification performance (Figures S3 - S5).

#### 2.4.2. Effects of image timing and geography

Seasonal variation in tree appearance and regional differences in genus composition (e.g., differing species) may influence genus-level classification rates. We analyzed image classification performance in relation to the month of GSV image capture and geography.

We grouped cities into four climate regions (Figure S6) using a principle component analysis (PCA) using 19 bioclimatic variables from WorldClim v2.1 at 2.5’ (∼4.5 km) resolution (https://worldclim.org/) that capture temperature and precipitation patterns relevant to tree distributions and phenology (Boucher-Lalonde et al., 2012; Fick and Hijmans, 2017). Bioclimatic variables were sampled at each city’s centroid using the raster R package (Hijmans et al., 2015). The first two PCA axes explained 77.6% of inter-city climate variation. Cities were then assigned to one of four regions based on their scores along these axes (Table 1, Figure S6).

We computed F1 scores for each genus by month and region combination using the 30% geographically independent test set (Figure S2). Because many genera lacked imagery across all month–region combinations, we restricted the analysis to genera represented in at least 50 month-region combinations (N = 17). This criterion increased temporal and spatial coverage for trend detection and reduced model failures due to missing data.

For the remaining 17 genera, we assessed the effects of month, region, and their interaction on classification performance using a two-way ANCOVA (Type II SS). The number of training images for each genus–month–region combination was included as a covariate to evaluate the influence of training data availability. Model assumptions were checked with standard diagnostics before inference. Pairwise comparisons of estimated marginal means for month and region used Tukey’s adjustment via the ‘emmeans’ R package (Lenth, 2025).

### 2.5 Generalizability of computer vision-based tree inventories

To evaluate generalizability, we applied our computer vision models across cities (Figure 1) and compared predicted trees to AutoArborist inventory records. Predicted and inventory trees were paired using a greedy matching approach, and we modeled the probability of a matched pair with binomial generalized linear mixed effects models (GLMMs; logit link).

#### 2.5.1 Sampling inventory trees

We restricted analysis to inventory trees located within the 30% geographic test split of each city (Figure S2). To capture local variability, each city was tiled into 100-m grid cells, and we retained only grids containing at least one tree. We then stratified grids by tree density using the 0.25, 0.5, 0.75 quantiles and drew a 5% random sample without replacement within each stratum. This design yielded 103,428 trees across 6,306 grids in 23 cities (Table S2).

For each sampled grid, we downloaded all available GSV images intersecting the grid and a 100-m outward buffer to reduce edge effects from missing imagery (Supplementary Methods 1.3). We then applied the computer vision models to these GSV images to detect, classify, and geolocate trees.

#### 2.5.2 Matching predictions to inventory trees

Predicted trees were linked to inventory trees using a greedy, one-to-one genus-matching approach. For each inventory tree, we computed a 20 meter circular buffer and identified all congeneric predicted trees within this area. Geodesic (great-circle) distances in meters were computed from WGS84 geographic coordinates for every inventory-prediction pair. We then iteratively selected the closest available pair, recorded the match, and removed both trees from future consideration, yielding a one-to-one assignment. Inventory trees remained unmatched if no congeneric predictions fell within their buffer or if all such predictions were already matched. We tested buffer radii from 5 to 30 meters in 5 meter increments and adopted 20 meters to balance matching performance with a local spatial scope (Figure S12).

#### 2.5.3 Effects of ecological, image coverage, and prior-knowledge on match success

We modeled the probability that an inventory tree matched a congeneric prediction using binomial GLMM. The full model was:

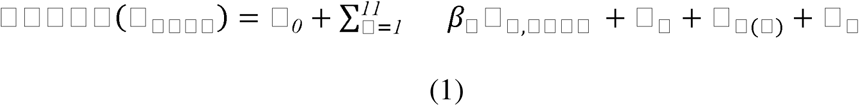

where □_□□□□_ is the probability that an inventory tree □ in a grid □ of a city □ belonging to a genus □, was correctly matched (1 = match; 0 = no match). □_0_ is the intercept, and □_□,□□□□_ are the 11 predictors. Random intercepts were specified as 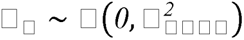 for city, 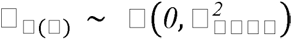 sampling grids nested within city, and 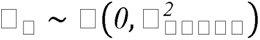 for genus.

We evaluated 11 fixed effects hypothesized to influence model generalizability: Ecological conditions (5): tree abundance, congeneric tree abundance, genus richness, nearest-neighbor (NN) tree distance (m), and temperature seasonality. These capture local tree density and spatial structure, which may influence matching outcomes by varying occlusion (tree abundance, NN tree distance), misclassification risk (genus richness), and match likelihood (congeneric abundance). Ecological predictors were computed within a 20 m buffer centered on each inventory tree, except seasonality, which extracted for each grid from the WorldClim v2.1 dataset (30” resolution, ∼1 km x 1 km; https://worldclim.org/) using the ‘terra’ R package (Hijmans, 2022).

Street-level image coverage (4): number of images, nearest image distance (m), modal capture year, and modal capture season (i.e., growing or dormant season; Supplementary Methods 1.4). These quantify the street-level information environment which may influence matching outcomes by affecting viewpoint availability and perspective diversity (number of images, nearest image distance), temporal recency and resolution (capture year), and visibility of diagnostic traits (capture season). Image coverage predictors were computed within a 20 m buffer centered on each inventory tree.

Prior knowledge bias (2): the number of AutoArborist training images for the tree’s genus, and the proportion originating from the focal city (the city-level count divided by the genus-level count). These predictors quantify the model’s genus- and location-specific familiarity, which impacts tree classification, and thus matching outcomes.

We included random intercepts for city and genus to account for data hierarchy, and a nested grid-within-city intercept to account for spatial autocorrelation. All numeric predictors were centered and scaled, and skewed predictors (all except temperature seasonality and capture year/season) were log-transformed to improve linearity on the logit scale.

We fit all 2,046 additive submodels of the full specification in R v.4.5.1 (R Core Team, 2025) using the ‘lme4’ (Bates et al., 2015) and ‘MuMIn’ (Bartoń, 2024). Models were ranked by the Bayesian Information Criterion (BIC) as a conservative choice for large samples (Schwarz, 1978). Diagnostics for the top-ranked (minimum BIC) model included residual plots, uniformity, dispersion, and outlier checks with ‘DHARMa’ (Hartig, 2024), variance inflation factors with ‘performance’ (Lüdecke et al., 2021), and spatial autocorrelation tests with ‘spdep’ packages (Bivand, 2022) (Supplementary Methods 1.5)

We assessed consistency in variable inclusion, effect sizes, and directions for the full model and all submodels with ΔBIC < 10. For the top model, we report marginal R^2^ (variance explained by fixed effects) and the conditional R^2^ (variance explained by fixed and random effects) using ‘performance’ package (Lüdecke et al., 2021). We decomposed random-effect variances to estimate the percentage of total variance attributable to city, grid, and genus. We visualized fixed-effect influences with partial effect plots, and deviations of city and genus levels from overall means were examined using best linear unbiased predictions (BLUPS).

## 3. Results

### 3.1 Tree detection and classification

#### 3.1.1 Object detection accurately identifies trees in street-level images

The object detection model identified trees with high precision (92.2%) and recall (82.4%) when evaluated using withheld testing images (Figure S7). Both high precision and recall values indicate a low proportion of false positive detections and that the model identified most trees present in GSV images.

#### 3.1.2 Classification performance and model interpretation

Classification performance varied widely among 100 genera (mean F1 score = 0.55; range: *Phoenix* [date palm], F1 = 0.88; *Carya* [hickory], F1 = 0.06; Table S3). In general, higher performance was correlated with a greater number of training images (*p < 0.001*; R^2^ = 0.50; Figure S8). However, the genus with the highest performance (*Phoenix* [date palm]; N = 3,790, F1 = 0.88) had considerably fewer training images than the most common class (*Acer* [maples]; N = 45,743; F1 = 0.75).

The Grad-CAMs provided insight into different visual features underlying correct genus classifications (Figure 2). For example, correctly classified *Citrus* trees emphasized the characteristic orange fruit (Figure 2C). For genera where fruiting or flowering were absent, structural features, such as canopy shape and branching patterns, were important for classification (e.g., for deciduous genera like *Acer* and *Betula*, and for palms like *Phoenix*) (Figure 2). Likewise, bark color and texture informed correct classifications. The reddish-brown, furrowed bark in conifers (*Pinus*, *Thuja*), and white-brown papery bark in *Betula* (birch) guided correct classifications (Figure 2). Across genera, Grad-CAMs revealed that classification relied on biologically meaningful and visually distinctive features.

**Figure 2.**
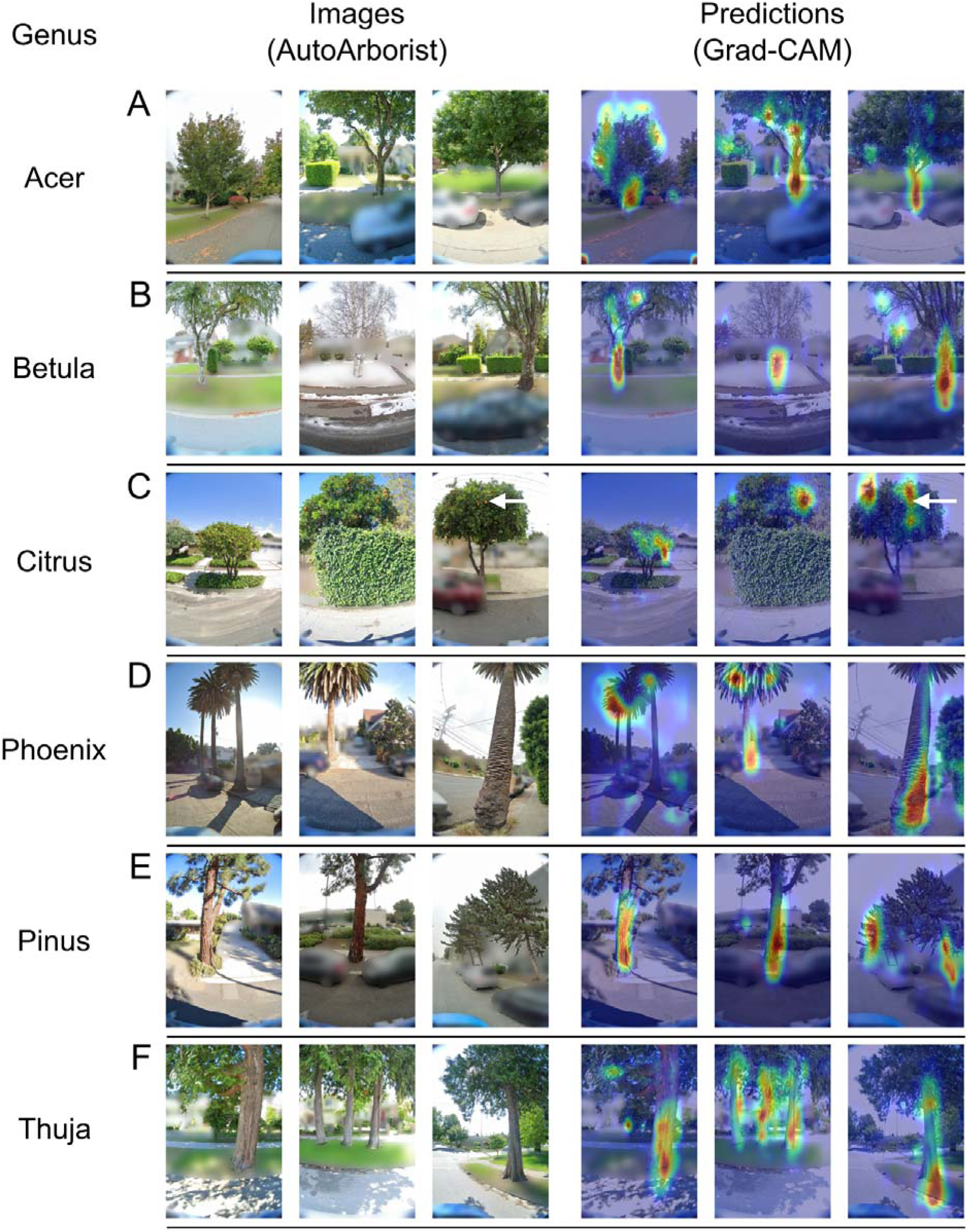
Representative gradient-weighted class activation maps (Grad-CAMs) for six tree genera classified by the convolutional neural network (CNN). Each row shows three AutoArborist images (left) and three corresponding Grad-CAMs (right) for one genus: (A) *Acer* (Maple), (B) *Betula* (Birch), (C) *Citrus* (Citrus), (D) *Phoenix* (Date Palm), (E) *Pinus* (Pine), and (F) *Thuja* (Arborvitae). Grad-CAMs highlight image regions most influential for the model’s predictions (red = higher importance; blue = lower importance). The CNN leveraged biologically meaningful and visually distinctive features, including reproductive traits (white arrow indicating fruit on *Citrus*, panel C), structural characteristics (e.g., canopy shape and branching in *Acer*, *Betula*, and *Phoenix*, panels A–D), and bark texture (e.g., reddish-brown furrowed bark in *Pinus* and *Thuja*, and papery white bark in *Betula*, panels B, E, F). These patterns show that the model captured diagnostic morphological features consistent with expert knowledge, supporting the interpretability of genus-level classifications.

### 3.2 Impact of diversity, image timing, and geography on genus classification performance

#### 3.2.1 Urban tree diversity affects performance among cities

Genus diversity within cities impacted overall classification performance. Cities with a higher Shannon diversity index had significantly lower performance than less-diverse cities (*p < 0.001*; Figure 3, Table S4). Likewise, cities with a greater genus richness had lower performance (*p = 0.045*; Figure S9, Table S4). The relative abundance of genera also affected accuracy (*p < 0.001;* Figure S9, Table S4). Cities where few genera are dominant (e.g., Sioux Falls, SD; evenness = 0.48) had higher performance than cities where no single genus dominates (e.g., Los Angeles, CA; evenness = 0.89) (Table S4).

**Figure 3.**
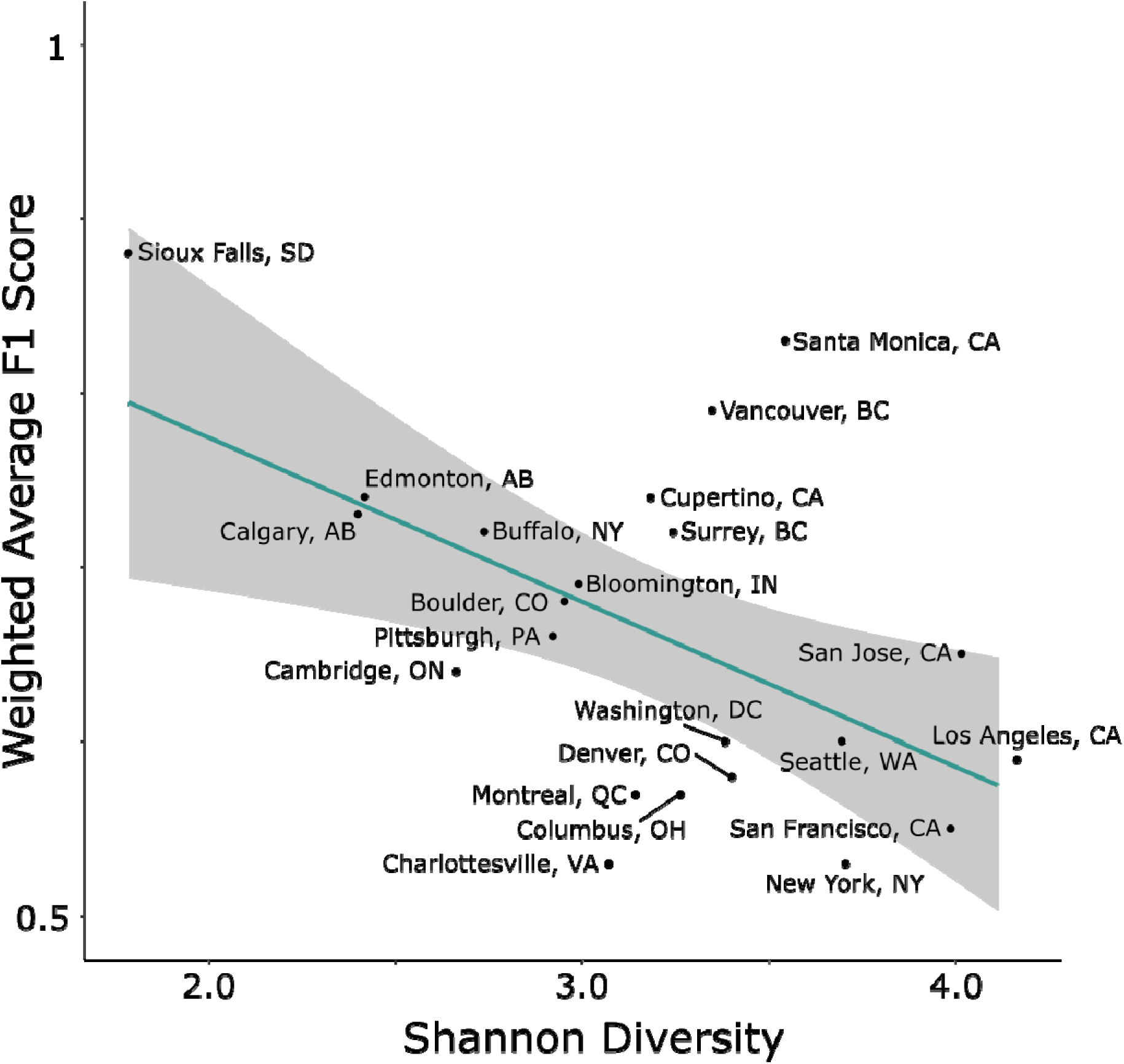
Relationship between the city-level Shannon diversity index and image classification performance across 23 cities. For each city, the Shannon diversity index was computed using from AutoArborist inventory records. We applied the image classification model to a 10% randomly withheld testing dataset for each city and evaluated performance by computing the weighted average F1 score across genera. The scatterplot illustrates the negative relationship between Shannon diversity and performance, where more diverse cities have lower weighted F1 scores. The teal line indicates the predicted values and shading denotes the 95% prediction interval.

#### 3.2.2 Geography and capture month affect performance

Genus classification performance varied by geographic region and, to a lesser extent, by the month of GSV image capture (Table S5). We detected regional differences in F1 score for the majority of tested genera, including *Acer* (maple), *Fraxinus* (ash), *Quercus* (oak), *Ulmus* (elm), *Prunus* (cherry/plum), *Tilia* (basswood), *Malus* (apple), *Platanus* (sycamore), *Magnolia* (magnolia), and *Crataegus* (hawthorn) (all *p < 0.01*; Table S5, Figure 4, Figure S10).

**Figure 4.**
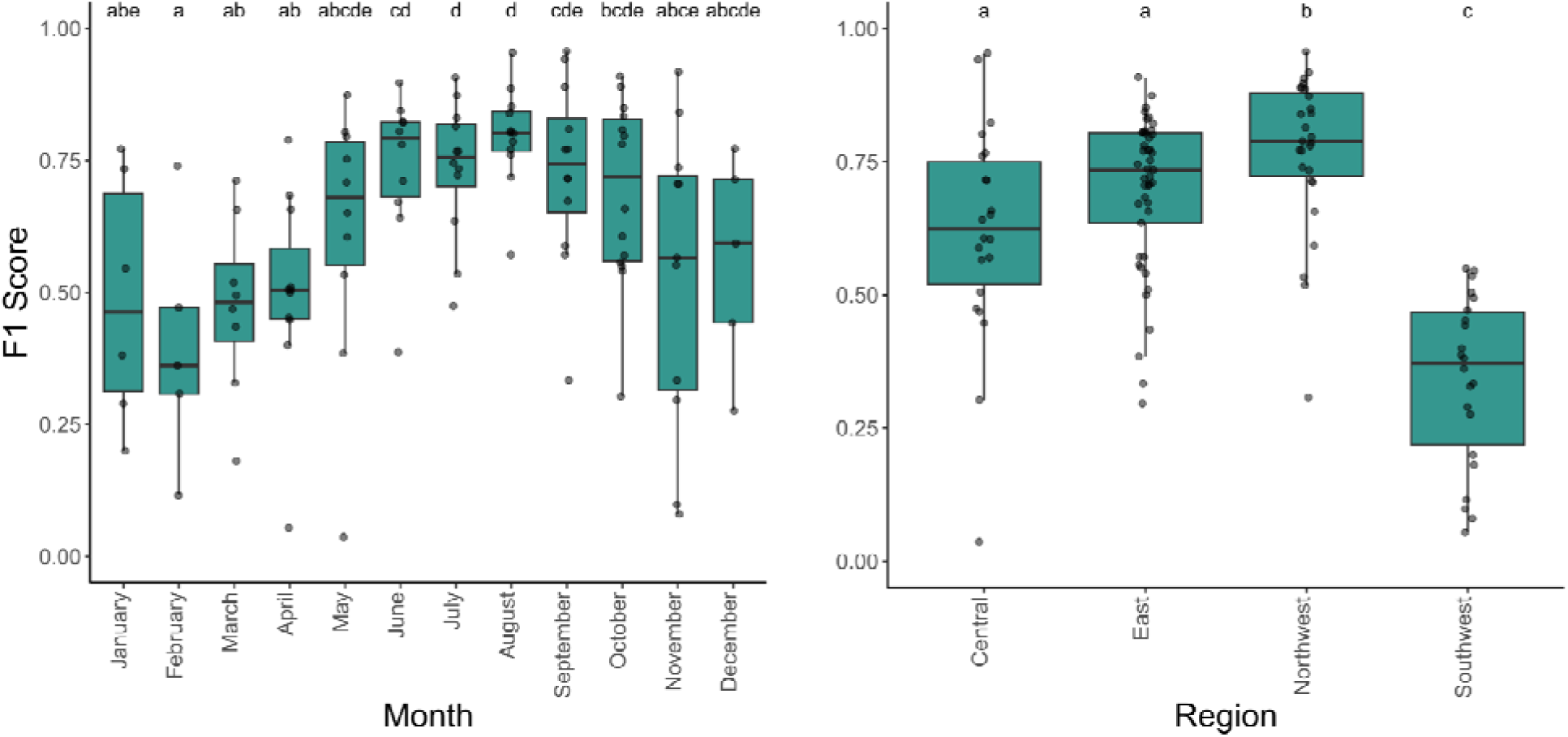
Maple (*Acer*) classification performance by month and climate-defined geographic region. Boxplots show F1-score distributions across months of image capture (left) and regions (right). Regions were derived by principal component analysis of bioclimatic variables, grouping cities by climate similarity. Each point represents the F1 score for testing images from a specific city-month combination within the 30% geographic testing split. Differences in performance were assessed with a two-way ANCOVA (Month, Region; Type II), followed by Tukey-adjusted pairwise comparisons of estimated marginal means; letters above boxplots denote significantly different groups.

Image capture month significantly influenced classification for the most common genus, *Acer* (*p < 0.001*; Table S5, Figure 4). Images from summer months (e.g., July - August) produced fewer false positives and false negatives than winter images (e.g., January - February) (Figure 4). This effect was independent of the monthly sample size and was consistent across regions (Table S5). For all other tested genera, monthly differences were not significant (*p > 0.05;* Table S5, Figure S11).

### 3.3 Generalizability of computer vision-based tree inventories

#### 3.3.1 Baseline matching rates across genera and cities

Across cities and genera, the computer vision-based inventory matched 67.1% of the inventory trees to predicted congeneric trees. Matching varied more across genera (mean = 50.9%, SD = 23.0%; Table S6) than across cities (mean = 67.4%, SD = 9.4%; Figure S12).

#### 3.3.2 Model Selection

We evaluated 2,046 candidate models to test how ecological, image coverage, and prior knowledge predictors affected generalizability. The best-supported model (BIC weight = 0.702) retained eight predictors from the full specification: four ecological (tree abundance, genus richness, congeneric abundance, nearest-neighbor tree distance), two image coverage (nearest image distance, number of images), and two prior-knowledge predictors (the number of AutoArborist training images for the tree’s genus, and the proportion originating from the focal city).

Two additional models (ΔBIC < 10) received limited support. These differed from the top model by either excluding genus richness or including the modal capture season as a fixed effect. The full model was strongly over-parameterized (ΔBIC = 30.77) (Figure 5A).

**Figure 5.**
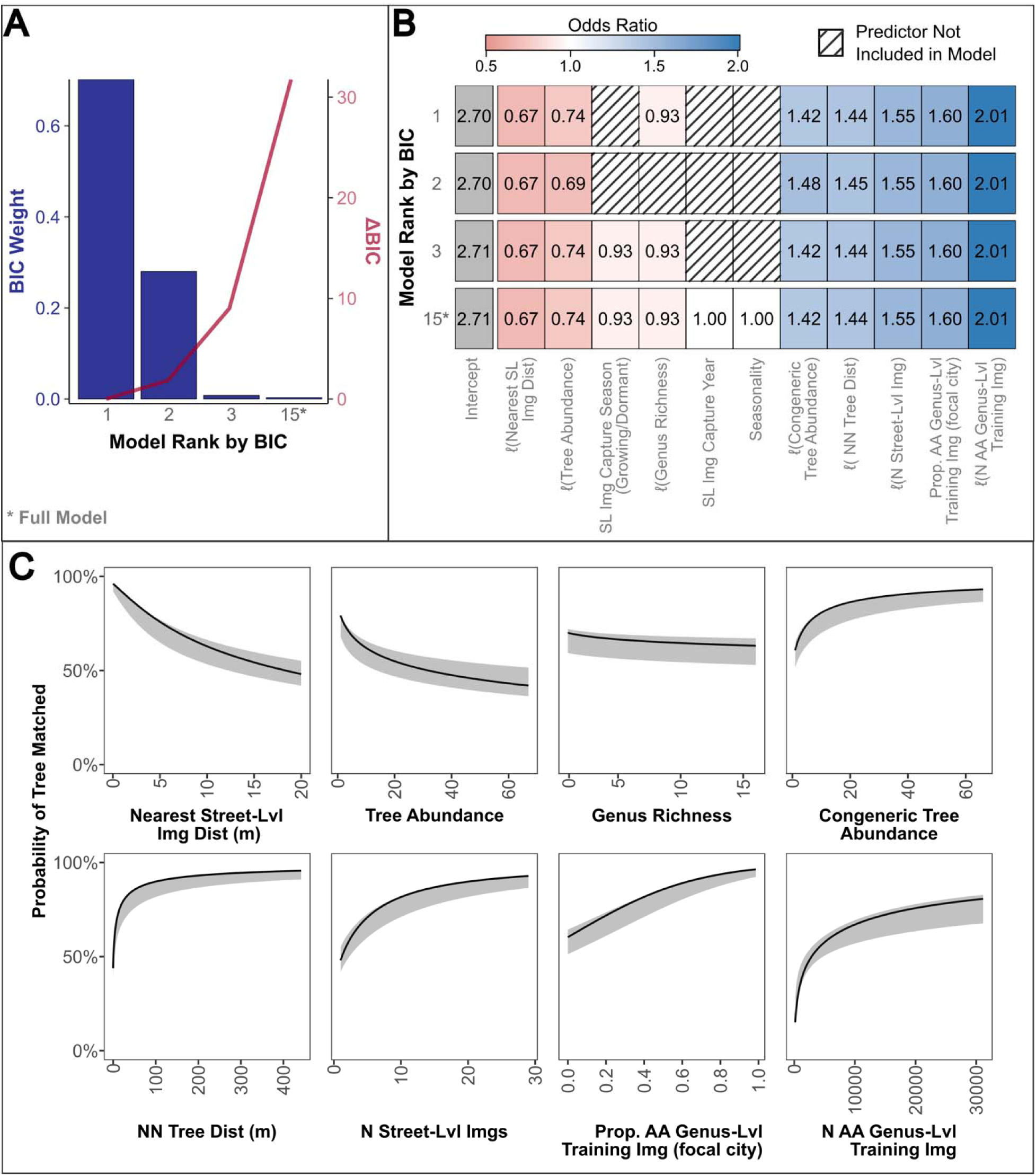
GLMMs evaluating how ecological context, street-level imagery coverage, and prior knowledge affect the probability that computer-vision predictions match AutoArborist inventory records from a geographically independent test set. Only the full model and submodels with a ΔBIC < 10 are shown. (A) Relative BIC weights (bars, left axis) and ΔBIC values (line, right axis) for each GLMM. (B) Heatmap of fixed-effect coefficients expressed as odds ratio; striped tiles denoting predictors excluded from a given model. Numeric predictors were transformed as [ℓ - log(x+1)]. (C) Partial effects showing predicted probabilities of matching across each predictor’s observed range, with all other predictors held at their mean. Solid lines and shaded ribbons denote model predictions and 95% confidence intervals, respectively. Predictor values are back-transformed to the original scale. Predictions are obtained from the inverse link function with bias correction to account for Jensen’s inequality. Abbreviations: SL = street-level; AA = AutoArborist.

Effect sizes for predictors shared across supported models (ΔBIC < 10) were highly consistent in magnitude and direction (mean SD of fixed-effect odds ratio estimates = 0.008; Figure 5B). Predictors unique to specific models exerted negligible influence on matching. Given the stability of effects and the strong support for the top model, all subsequent inferences are based on that model.

#### 3.3.3 Model diagnostics and goodness of fit

Diagnostics for the top model indicated that assumptions were met: no overdispersion, non-uniform residuals, influential outliers, or spatial autocorrelation (Figures S13–S14). Fixed effects explained 20.5% of the variance in matching (marginal R^²^), increasing to 46.7% when random effects were included (conditional R^²^). Random effect variance was dominated by genus (Var = 0.78, SD = 0.89), with a smaller contribution from city (Var = 0.46, SD = 0.46). These results are consistent with variance partitioning estimates, which attribute 12.7% of the total variance to genus and 3.5% to city (Table 2).

**Table 2.** Top GLMM (selected by minimum BIC) assessing how ecological, street-level image coverage, and prior-knowledge influence the probability that computer-vision predictions match AutoArborist inventory records in a geographically independent test set. Numeric predictors were transformed as [ℓ - log(x+1)]. Predictors associated with lower odds: nearest image distance, tree abundance, and genus richness. Predictors associated with higher odds: congeneric abundance, nearest-neighbor (NN) tree distance (m), the number of training images from the tree’s genus, and the proportion of training images.

| Fixed Effects |  |  |  |  |  |
| --- | --- | --- | --- | --- | --- |
| Parameter | OR | SE | 95% CI | Wald z | Sig. |
| Intercept | 2.70 | 0.48 | [1.91, 3.82] | 5.59 | *** |
| ℓ(SL Img Distance) | 0.67 | 0.01 | [0.65, 0.68] | -37.97 | *** |
| ℓ(Tree Abudance) | 0.74 | 0.02 | [0.71, 0.77] | -13.10 | *** |
| ℓ(Genus Richness) | 0.93 | 0.02 | [0.90, 0.97] | -3.67 | *** |
| ℓ(Congeneric Abundance) | 1.42 | 0.02 | [1.37, 1.46] | 20.45 | *** |
| ℓ(NN Dist) | 1.44 | 0.02 | [1.41, 1.48] | 28.75 | *** |
| ℓ(N SL Img) | 1.55 | 0.02 | [1.51, 1.58] | 39.01 | *** |
| Prop. AA Genus-Lvl Training Img (focal city) | 1.60 | 0.03 | [1.53, 1.67] | 22.11 | *** |
| ℓ(AA Genus-Level Training Img) | 2.01 | 0.16 | [1.71, 2.35] | 8.63 | *** |
| Random (Intercept) Effects |  |  |  |  |  |
| Grouping Factor | N | Var. | SD | % Total Var. |  |
| Grid (City) | 5864 | 0.62 | 0.78 | 12.7 |  |
| Genus | 100 | 0.78 | 0.89 | 10.0 |  |
| City | 23 | 0.46 | 0.46 | 3.5 |  |
| Residual | - | - | - | 73.8 |  |
| Model Fit |  |  |  |  |  |
| Metric | Value |  |  |  |  |
| N | 90689 |  |  |  |  |
| Degrees of Freedom | 12 |  |  |  |  |
| Marginal R² | 0.205 |  |  |  |  |
| Conditional R² | 0.467 |  |  |  |  |

**Notes:** OR = odds ratio; SE = standard error; CI = confidence interval; Sig. = Significant; NS = Not significant; $\ell$ = $\log(x + 1)$ ; AA = AutoArborist; SL = Street-Level
\* $p < 0.05$ ; \*\* $p < 0.01$ ; \*\*\* $p < 0.001$

#### 3.3.4 Influence of ecological, street-level image coverage, and prior-knowledge predictors

All fixed effects retained in the top model significantly affected the probability of correctly matching inventory trees to predicted trees (Table 2). Prior-knowledge exerted the strongest effects. A multiplicative increase in the number of genus-level training images (log-transformed) more than doubled the odds of a match (OR = 2.01, 95% CI: 1.71–2.35, *p < 0.001*) and a higher proportion of training images originating from the focal city also increased success (OR = 1.60, 95% CI: 1.53–1.67, *p < 0.001*).

Local image coverage had comparably strong effects. More images (log-transformed) improved matching (OR = 1.55, 95% CI: 1.51-1.58, *p < 0.001*), while greater distance to the nearest image (log-transformed) reduced matching (OR = 0.67, 95% CI: 0.67–0.68, *p < 0.001*).

Local ecological context showed mixed effects. Greater nearest-neighbor tree distances (log-transformed; OR = 1.44, 95% CI: 1.41–1.48, *p < 0.001*) and higher congeneric abundance (log-transformed; OR = 1.42, 95% CI 1.37–1.46, *p < 0.001*) increased matching odds, whereas higher tree abundance (log-transformed; OR = 0.74, 95% CI: 0.71–0.77, *p < 0.001*) and greater genus richness (log-transformed; 0.93, 95% CI: 0.90–0.97, *p < 0.001*) decreased matching (Figure 5B–5C). Together, matching is maximized when the tree’s genus is well represented in the training dataset, particularly with images from similar geographic contexts, when image coverage is dense, and when the local tree composition and spatial arrangement are simpler.

#### 3.3.5 Genus- and city-level deviations

Genus-level best linear unbiased predictions (BLUPs) showed consistent overperformance and underperformance after accounting for fixed effects (Figure S15A). Among the 100 genera, 29 genera overperformed and 21 underperformed (defined as BLUP 95% CIs not overlapping the overall mean). Largest positive deviations were observed for *Ailanthus* (tree-of-heaven; BLUP ± 95% CI: 1.99 ± 0.39), *Melaleuca* (paperbark; 1.84 ± 0.37), *Abies* (fir; 1.56 ± 0.54), *Pseudotsuga* (douglas fir; 1.35 ± 0.38), and *Sequoia* (coast redwood; 1.22 ± 0.58). The strongest negative deviations occurred for *Ostrya* (hop hornbeam; –1.75 ± 0.76), *Cornus* (dogwood; –1.63 ± 0.26), *Chamaecyparis* (false cypress; –1.52 ± 0.67), *Syringa* (lilac; –1.44 ± 0.22), and *Citrus* (citrus; –1.33 ± 0.72).

City-level BLUPs (Figure S15B) were more constrained, consistent with their smaller share of explained variance (3.5%; Table 2). Three cities overperformed (Vancouver = 1.20 ± 0.21; Charlottesville = 0.85 ± 0.54; Buffalo = 0.56 ± 0.28), and five underperformed (Kitchener = –0.89 ± 0.36; Los Angeles = –0.46 ± 0.16, Washington, DC = –0.35 ± 0.18, San Francisco = –0.26 ± 0.18, Edmonton = –0.22 ± 0.19). Overall, genus identity remains the primary source of residual variation in matching performance, whereas city-level deviations are fewer and modest in magnitude.

## 4. Discussion

Urban tree inventories underpin management of ecosystem services and responses to disturbances such as pest or disease. Although existing remote sensing methods accurately map trees within individual cities, their ability to generalize across variable imaging conditions, species compositions, and climates remains uncertain. We addressed this knowledge gap by training computer vision models on GSV and iNaturalist images to detect, classify, and map 100 tree genera across 23 North American cities. Object detection and depth estimation reliably identified and geolocated trees, but genus classification performance was more variable. Accuracy declined with shifts in genus diversity and geography but improved under leaf-on conditions. Grad-CAM indicated that classification leveraged biologically interpretable traits (e.g., canopy structure, bark texture, and fruit). In geographically independent evaluations, predictions matched 67.1% of AutoArborist inventory records. Variation in matching success was driven primarily by differences among genera, training data availability, and local ecological and image context rather than city identity. Collectively, these results show that street-level imagery coupled with computer vision offers a scalable complement to traditional inventories.

### 4.1. Opportunities and challenges for scaling urban tree inventories

Street-level imagery provides a promising alternative to aerial or satellite data for urban forest monitoring. While prior applications were city-specific (Branson et al., 2018; Capecchi et al., 2023; Choi et al., 2022; Liu et al., 2023; Velasquez-Camacho et al., 2024; Wegner et al., 2016), our analysis across 23 cities enabled a broader assessment of model generalizability. The models achieved high precision and recall (>0.7) for eight common U.S. genera (*Acer, Fraxinus, Quercus, Gleditsia, Ulmus, Tilia, Pyrus,* and *Platanus)* (Table S3). Given these taxa represent ∼45% of public street trees in the U.S., (Ma et al. 2020), this approach has operational value for many municipalities. For instance, our approach performed strongly in Sioux Falls, SD, because ∼82% of trees belong to these genera (Figure 3).

Performance declined in large, biodiverse cities such as Los Angeles, New York, and Denver, likely reflecting visual similarity among co-occurring genera (low inter-class variability) and numerous species and cultivars (high intra-class variability). This pattern mirrors findings from field and remote sensing studies showing reduced classification performance in taxonomically diverse areas (Berland et al., 2019; Marconi et al., 2022; Weinstein et al., 2023). Although we classified at the genus level, future work could predict across multiple taxonomic levels and leverage phylogenetic relationships to improve accuracy (Gillespie et al., 2024).

### 4.2 Interpreting model predictions with trait-based evidence

Model performance varied by geography and seasonality, likely reflecting regional differences in tree composition, management practices, and contextual cues such as street layouts (Beery et al., 2022). Image timing also strongly influenced classification. For *Acer*, leaf-on months (July–August) reduced both false positives and negatives compared with winter imagery, independent of training sample size (Table S5, Figure 4).

Grad-CAM visualizations clarified how models distinguished genera. Some taxa were identified by fruits or flowers, while others relied on subtler traits like canopy architecture and bark texture. These findings align with prior work showing that CNNs can reliably identify species using tree bark imagery (Kim et al., 2022) and suggest that the visual cues driving classification shift seasonally. Bark provides consistent information year-round, whereas flowering and fruiting periods offer brief but highly informative windows for genus identification. Feature salience may also vary with tree age. Grad-CAMs highlighted age-related cues such as trunk buttressing and coarse bark patterning that are pronounced in mature trees but less apparent in young saplings. For younger trees, foliage traits may therefore provide more informative signals for classification. These results point to opportunities for optimizing image acquisition by aligning data collection with periods and contexts when locally diagnostic traits are most visible.

### 4.3 Factors shaping generalizability across cities

Generalizability was driven by training data volume, local ecological structure, and street-level image coverage. Genus-level image availability was the strongest predictor of matching success: > 3,000 training images per genus were typically required to exceed a 50% match probability. Incorporating city-specific training images further improved performance: matching rose from 61% to 75% as the share of focal-city training images increased from 1% to 25% (Figure 5B–C). Denser stands reduced match success, consistent with occlusion effects by neighboring trees: match probability fell below 50% when > 30 trees occurred within 20 meters of the inventory tree and a minimum spacing of ∼1 m was needed to achieve a 50% match probability, increasing to 75% at 15 m (Figure 5B–C). Image density and camera-to-tree distance also mattered: match probability rose from 50% to 63% as available viewpoints increased from one to three, but dropped below 50% when the nearest image distance exceeded 18 m

These quantitative relationships extend qualitative observations in previous studies (e.g., Lumnitz et al., 2021; Wegner et al., 2016). Operationally, computer vision inventories will perform best in well-imaged areas, with leaf-on imagery that shows discriminative features, and in structurally simple stands (i.e., limited crowding and occlusion). Targeted expansion of training images, particularly for underrepresented genera with examples from the target geography, can further improve reliability of these approaches.

### 4.4. Model limitations and implications for urban forestry

Despite strong performance, several limitations remain. First, public tree inventories may be incomplete or outdated; some apparent model errors may reflect inaccuracies in the reference data. Second, our matching rates are conservative because predictions were compared only to public inventories, and private trees could not be assessed. Third, our 20-m matching radius introduces uncertainty by potentially conflating nearby public and private trees. A stricter 15-meter radius reduced overall agreement to 60.0%, but the broader statistical patterns remained unchanged (Figure S12). Last, most inventories lacked tree-health attributes, limiting our ability to distinguish between healthy, stressed, and diseased trees (but see Kagan et al., 2021).

Integrating multiple image sources and spatial context could reduce errors. National Agricultural Imagery Program imagery (NAIP) provides high-resolution, leaf-on coverage suitable for urban tree mapping (Erker et al., 2019; Rice et al., 2025; Ventura et al., 2024). Fusing NAIP with street-level imagery can improve classification rates by providing complementary canopy and contextual information (Hartling et al., 2019; Wang et al., 2025). Conversely, others report marginal gains from incorporating high-resolution imagery (Beery et al., 2022). Adding geographic context during training could also improve fine-grained classification. Given that species distributions are governed by environmental gradients (Boucher-Lalonde et al., 2012), incorporating location can constrain predictions to ecologically plausible taxa (Aodha et al., 2019; Cole et al., 2023).

Understanding where and when models generalize remains a core challenge, particularly under class imbalance (Zhou et al., 2023; Beery et al., 2018). The AutoArborist dataset (Beery et al., 2022) exemplifies this challenge by reflecting the long-tailed distribution of urban forests, where few genera dominate and most are rare. Our results demonstrate computer vision applied to street-level imagery has scalable applications for urban forestry management despite this imbalance. Extending these methods to peri-urban and rural landscapes could support biodiversity assessments and invasive species monitoring (Kotowska et al., 2021). Where street-level imagery is sparse, fusing street-level and aerial data offers a practical strategy to fill coverage gaps.

## 5. Conclusions

Computer vision models trained on publicly available street-level imagery can accurately detect, classify, and geolocate urban street trees across diverse North American cities. Where classification rates declined, namely in areas with high genus richness or off-season imagery, we identified failure patterns, such as inter-class confusion (between-genus look-alikes) and intra-class variability (visual variation within a genus) that mirror challenges faced by community observers and arborists. By quantifying where and when performance deteriorates and identifying practical levers for improvement (e.g., leaf-on acquisition, denser image coverage, targeted expansion of training images), we establish a framework for systematically improving future urban tree mapping. Developing scalable, generalizable models for tree inventories will be essential for monitoring and safeguarding ecosystem services amid growing environmental pressures.

## CRediT authorship contribution statement

**Thomas Lake**: Conceptualization, Data curation, Formal analysis, Investigation, Methodology, Resources, Software, Project administration, Validation, Visualization, Writing – original draft, Writing – review & editing. **Brittany Laginhas**: Conceptualization, Data curation, Formal analysis, Investigation, Methodology, Resources, Software, Project administration, Validation, Visualization, Writing – original draft, Writing – review & editing. **Ross Meentemeyer**: Conceptualization, Funding acquisition, Project administration, Supervision, Writing – review & editing. **Chris Jones**: Conceptualization, Data curation, Project administration, Resources, Supervision, Validation, Writing – original draft, Writing – review & editing. **Brennen Farrell**: Data curation, Formal analysis, Investigation.

## Acknowledgements

This work was made possible by a data sharing agreement (https://google.github.io/auto-arborist). We are grateful to the community scientists who collected images used in the study. We acknowledge the computing resources provided by North Carolina State University’s High Performance Computing Services Core Facility (RRID: SCR_022168).

## Data accessibility

Code and models are available on Github (https://github.com/ncsu-landscape-dynamics/gsv_host_detector) and will be made publicly available upon acceptance.

