## Supplementary Materials for "Continental-scale computer vision models reveal generalizable patterns and pitfalls for urban tree inventories with street-view images"

**AUTHORS**: Thomas A. Lake^a, † *^
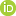
, Brit B. Laginhas^a, †^
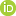
, Brennen T. Farrell^a^
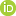
<https://orcid.org/0009-0004-4794-5906>, Ross K. Meentemeyer^a^
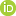
, and Chris M. Jones^a*^
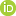

^a^North Carolina State University, Center for Geospatial Analytics, 5112 Jordan Hall, 2800 Faucette Dr, Raleigh, NC 27695

*Corresponding authors at: North Carolina State University, Center for Geospatial Analytics, 5112 Jordan Hall, 2800 Faucette Dr, Raleigh, NC 27695

^†^Co-first authors

Supplementary Methods

SM 1.1 Tree detection model

SM 1.1.1 Model architecture

SM 1.1.2 Model training

SM 1.1.3 Model evaluation

SM 1.2 Tree classification model

SM 1.2.1 Model architecture

SM 1.2.2 Model training

SM 1.2.3 Model evaluation

SM 1.3 Downloading street-level images

SM 1.4 Classifying the modal capture month into growing vs. dormant season

SM 1.5 Testing for spatial autocorrelation in tree-level matching among cities

Supplementary Figures

Figure S1. Examples of AutoArborist images, iNaturalist images, and filtered iNaturalist images with Contrastive Language-Image Pre-Training (CLIP) deep learning model.

Figure S2. Location of Google Street View imagery included in the AutoArborist dataset for 23 study cities with images divided by geographic splits.

Figure S3. Diagnostic plots for regression models on genus Shannon diversity and classification performance demonstrating the city of Kitchener, ON as an outlier.

Figure S4. Diagnostic plots for regression models on genus richness and classification performance demonstrating the city of Kitchener, ON as an outlier.

Figure S5. Diagnostic plots for regression models on genus evenness and classification performance demonstrating the city of Kitchener, ON as an outlier.

Figure S6. Map of 23 study cities and PCA biplot using WorldClim climate variables, grouping cities into four geographic regions.

Figure S7. Precision and recall performance curve for the YOLOv5x tree detection model evaluated at the Intersection over Union (IOU) threshold of 0.5.

Figure S8. Correlation between the number of training images used in the genus classification model and testing performance, as measured by F1 score.

Figure S9. Effects of genus richness and evenness on classification performance across cities.

Figure S10. Differences in genus classification performance among regions.

Figure S11. Differences in genus classification performance among months.

Figure S12. The proportion of inventory trees matched with at least one street-level image within the search radius, tested across radii of 5 m to 30 m, is shown overall and by city.

Figure S13. Residual diagnostics for the top binomial GLMM.

Figure S14. Correlogram of global Moran’s *I* for residuals from the top generalized linear mixed model (GLMM) with and without nested sampling grid random intercept.

Figure S15. City- and genus-level BLUPs from the top binomial GLMM showing variation in baseline tree matching probabilities with 95% confidence intervals.

Supplementary Tables

Table S1. Selected 100 tree genera examined in the study, and the number of training, validation, and testing images and sources used in the tree classification model.

Table S2. Counts of grids, genera, and inventory trees per city and totals used in the GLMM analysis.

Table S3. Performance metrics for image classification model of 100 tree genera.

Table S4. Beta-regression models between weighted average F1 score and diversity metrics across study cities.

Table S5. ANOVA tables for classification performance across genera by month, geography, and number of training images.

Table S6. Percent of inventory trees matched with at least one Google Street View image within the 20-m search radius by genus.

Supplementary References

**Supplementary Methods**

SM 1.1 Tree detection model

SM 1.1.1 Tree detection model architecture

To identify trees in street-level images, we trained a supervised deep learning model based on the YOLO (You Only Look Once) object detection architecture. The original YOLO model (YOLOv1; Redmon et al., 2016) is a one-stage object detection method that functions by dividing an input image into multiple regions (*S x S* grids) and predicting bounding boxes (*B*) for objects detected in each region. Each predicted bounding box consists of five values: the confidence score (representing how confident the model is that the box contains an object), the coordinates of the center of the box in the image (*b_x_, b_y_*), and the height and width of the box (*b_h_, b_w_*) relative to the full image (Terven and Cordova-Esparza 2023).

Since the development of the original YOLOv1 model, several iterations of the algorithm have improved the speed, accuracy, and ability to recognize multi-scale objects (Terven and Cordova-Esparza, 2023). We selected the YOLOv5x model for detecting trees in GSV images (Jocher et al., 2022; Redmon et al., 2016). Previous versions of the YOLO architecture (e.g., YOLOv3) have also previously shown success in detecting trees in street-view images (Choi et al., 2022; Liu et al., 2023).

The YOLOv5x model contains three main components: the backbone, neck, and head.

The backbone is responsible for extracting high-level features, such as edges, shapes, and textures, from input images through a convolutional neural network (CNN). The core component of CNNs is convolutional layers that apply small filters, or kernels, to local regions of an image. By performing this operation across the entire image, the model transforms the original image into feature maps, capturing patterns such as edges and shapes that are important for object detection. Here, the backbone was pre-trained on a large-scale image classification dataset (ImageNet) to facilitate feature extraction. The backbone uses multiple convolutional layers to extract hierarchical features at different scales, with lower-level features (e.g., edges, shapes, and textures) extracted in the earlier layers and higher-level features (e.g., object parts and semantic information) extracted in subsequent layers. Through training, the model learns the optimal parameters for each convolutional filter that result in accurate object detection.

The neck is an intermediate component that connects the backbone to the head. The neck is responsible for aggregating and refining features extracted from the backbone, and focuses on enhancing the spatial and semantic information across different scales. By processing features at multiple scales, the neck strengthens the model's ability to detect objects of various scales and sizes.

The head is the final component of the object detector; it is responsible for making predictions based on the features provided by the backbone and neck. Specifically, the head consists of several task-specific sub-networks that perform both object localization and classification. The object localization portion of the head generates predictions for each object candidate. Each predicted object is represented by a bounding box, class label (here, all labels are ‘tree’), and confidence score (ranging from 0 to 1), representing the model’s certainty in the prediction. The head also performs non-maximum suppression (NMS), which filters out overlapping predictions and retains only the most confident detections. As a result, we trained the YOLOv5x model to extract features and detect all trees in Google Street View (GSV) images.

SM 1.1.2 Model training

To train the object detection model, we used images of trees annotated with bounding boxes from the AutoArborist dataset (Beery et al., 2022). We randomly sampled 750,000 images of trees with annotated bounding boxes for model training. We withheld 75,000 images with annotations for a validation dataset to evaluate and fine-tune the models’ parameters during training. We reserved an additional 75,000 annotated images as an independent testing dataset to evaluate model performance.

Training the object detection model comprises two main steps: the forward pass and the backward pass:

Initially, the YOLOv5x models’ parameters (i.e., weights and biases) were initialized with pre-trained weights from ImageNet to improve feature extraction in the backbone. During the forward pass, samples of training data (in this case, 64 images of trees with annotated bounding boxes from AutoArborist) are fed through the model. As the images passed through each layer of the network, various computations were performed via convolutions using the model’s current parameters. The objective is to predict bounding boxes that are as close as possible to the true annotations.

The discrepancy between the model's predictions and the true bounding boxes is calculated as the training loss using a loss function. The training loss for the YOLOv5x model combines three primary components: objectness loss, classification loss, and box loss. Objectness loss is a measure of how well the model performs at predicting an object irrespective of the class. Box loss measures how close the predicted bounding box aligns with the true bounding box regardless of the class. Classification loss measures how well each bounding box classifies a detected object. Because our goal was to build an object detection model that identified all cases of a single class (i.e., ‘tree’) across all cities, the objectness and box loss played a more significant role than classification loss.

Once the loss is computed from the training data, an optimization algorithm determines how to adjust the model's parameters to improve its performance (i.e., reduce the loss). This is done during the backward pass. During this stage, the model calculates gradients, which represent the directions and magnitudes of change required for each parameter to minimize the loss. The YOLOv5x model used the stochastic gradient descent (SGD) optimizer with a learning rate of 0.01 and a momentum of 0.97 to update model parameters after each backward pass.

By repeating the forward and backward passes over multiple epochs—complete passes over the training dataset—the model progressively learns the intricate patterns in the data, allowing it to make accurate predictions when presented with new, unseen testing data. Throughout the training process, the validation dataset helps in tuning model parameters to prevent overfitting. After each epoch, the loss is also computed against the entire validation dataset (i.e., the validation loss). Validation loss is crucial because it helps assess how well the model generalizes to new, unseen data. Ideally, the validation loss should also decrease as training progresses. However, if it starts to increase significantly while the training loss continues to decrease, this indicates that the model might be overfitting to the training data. As such, we trained the YOLOv5x model for 25 epochs and monitored both the training and validation loss to limit overfitting.

To further improve model generalization to unseen images, we applied image augmentations during training. Image augmentations included random horizontal flipping (probability = 0.5), rescaling (±60%), rotation (±5°), and shearing (±1°). These transformations simulate potential variations in the appearance of trees in urban environments, helping the model generalize across different scales, orientations, and lighting conditions.

SM 1.1.3 Model evaluation

We evaluated the performance of the object detection model by calculating the number of true positive (TP), false positive (FP), and false negative (FN) detections. Performance was assessed at an Intersection over Union (IoU) threshold of 0.5. IoU is a metric that quantifies the overlap between a predicted bounding box and the ground-truth (annotated) bounding box. An IoU of 1 indicates a perfect overlap, whereas an IoU of 0 indicates no overlap.

We calculated precision as the ratio of true positive detections (TP) to the total number of predicted detections (TP + FP), which reflects the model’s ability to minimize false positives.

Precision = TP / (TP + FP).

We calculated recall as the ratio of true positive detections to the total number of ground-truth annotations (TP + FN), which measures the model’s ability to correctly detect objects.

Recall = TP / (TP + FN).

SM 1.2 Tree classification model

SM 1.2.1 Model architecture

To classify tree genera detected in street-view images, we developed a convolutional neural network (CNN) using the EfficientNetV2-S architecture (Tan and Le, 2021). The EfficientNetV2-S architecture uses a series of convolutional layers to extract high-level patterns, such as edges, textures, and shapes, from images that enable the model to distinguish types of images. CNNs learn parameters for convolutional layers that best extract discriminative features from the data. Images pass through multiple convolutional layers, allowing the CNN to learn combinations of features that lead to accurate image classification. The CNN captures hierarchical features at different scales, with lower-level features (e.g., edges and textures) extracted in the earlier layers and higher-level features (e.g., object parts and semantic information) extracted in subsequent layers. Through training, the model learns the optimal parameters for each convolutional layer that lead to the most accurate image classification.

The final layer of the image classifier transforms extracted features into a vector of predicted probabilities for each genus. We used a Softmax layer to normalize the vector of predicted probabilities on the scale of [0,1]. These predicted probabilities sum to one and can be interpreted as the likelihood of the genus in an image. We used an Argmax layer to select the genus with the highest probability as the class prediction.

SM 1.2.2 Model training

To train the image classification model, we used images of 100 tree genera from AutoArborist and iNaturalist (Table S1). Each image is labeled with a single genus. We randomly divided our dataset into training, validation, and testing sets based on an 80/10/10 split. Validation data was used to evaluate and fine-tune the model’s parameters during training, and testing data was withheld from model training and used as an independent set to evaluate performance.

Training the image classification model is similar to the object detection model and consists of two main steps: the forward pass and the backward pass:

We initialized the model’s parameters (i.e., weights and biases) with pre-trained weights from ImageNet. During the forward pass, samples of the input data (in this case, 64 images of tree genera from AutoArborist and iNaturalist) are fed through the model. As the data moves through each layer of the network, various computations are performed through a series of convolution layers using the model’s current parameters. The goal is to classify the actual tree genus. The discrepancy between the classification and the true genus is calculated as the training loss using a loss function. We used the categorical cross-entropy loss function to measure the difference between predicted labels and true labels for each class in a multi-class classification problem.

Once the loss is computed from the training data, an optimization algorithm determines how to adjust the model's parameters to improve its performance (i.e., reduce the loss). This is done during the backwards pass. During this stage, the model calculates gradients, which are essentially directions and magnitudes of change needed for each parameter to minimize the loss. We used the Adam optimizer (Kingma and Ba, 2014), a variant of stochastic gradient descent, to update the model’s parameters. Adam is a popular optimizer in deep learning because it incorporates adaptive learning rates and momentum, making the optimization process more efficient by speeding up convergence and stabilizing parameter updates.

By repeating the forward and backward passes over multiple epochs—complete passes over the training dataset—the model progressively learns to classify tree genera and make accurate predictions when presented with new, unseen testing data. Throughout the training process, the validation dataset helps in tuning model parameters to prevent overfitting. After each epoch, the loss is also computed against the entire validation dataset (i.e., the validation loss). Validation loss is crucial because it helps assess how well the model generalizes to new, unseen data. Ideally, the validation loss should also decrease as training progresses. However, if it starts to increase while the training loss continues to decrease, this indicates that the model might be overfitting to the training data. As such, we trained the image classification model for 100 epochs and monitored both the training and validation loss to prevent overfitting.

Prior to training, we normalized images on the scale of [0,1] using the mean and standard deviation of the red, green, and blue channels, respectively. During training, we applied image augmentations to enhance model generalization. For all images, we first used center cropping (768 x 768px) to standardize input image sizes and focus on the center of the image. We then applied random resize cropping (512px x 512px) to sample image portions at different scales (0.08 - 1.0) and aspect ratios (0.75 - 1.33). Additional augmentations included random horizontal flipping (probability = 0.5), color jittering (brightness, contrast, saturation, hue adjusted by factors of ±0.1), and random affine transformations (±5° rotations).

During training, we applied label smoothing (alpha = 0.05) to the categorical cross-entropy loss function as an additional regularization technique to account for images with multiple genera. Because images are labelled with a single genus (e.g., *Acer*), but in practice may have multiple genera present, we applied label smoothing to reduce the certainty that the class is solely present in an image (Müller et al., 2019).

We used a learning rate scheduler to reduce the initial learning rate of 0.001 by a factor of 10 if the validation loss increased for 5 consecutive epochs. This strategy promotes stable initial training and allows for efficient model convergence. We applied early stopping to terminate model training when the validation loss increased for 10 consecutive epochs, which indicates overfitting. We developed the image classification model with Python v3.10.13 and PyTorch v2.1.2 (Abadi et al. 2016; Paszke et al. 2017).

SM 1.2.3 Model evaluation

We assessed model performance for each genus with several evaluation metrics that are based on a class-wise confusion matrix from the Argmax output. The confusion matrix contains the number of correctly predicted true positives (TP), incorrectly predicted false positives (FP), correctly predicted true negatives (TN), and incorrectly predicted false negatives (FN) for all images in the withheld testing dataset.

The evaluation metrics we calculated were: Precision, Recall, and F1-score.

We calculated class-wise precision, or the proportion of all identified positives that are correct.

Precision = TP / (TP + FP).

We calculated class-wise recall, or the ability to predict all true positives (i.e., True Positive Rate (TPR)).

Recall = TP / (TP + FN).

We calculated the F1-score per class. F1 is derived from precision and recall. F1 scores are used when both precision and recall are important factors for model evaluation.

F1-score = (2 * precision * recall) / (precision + recall)

We additionally calculated the weighted average F1-score, which accounts for class imbalance by giving more weight to classes with more samples. This metric is calculated as the weighted mean of individual class F1-scores, where the weights are the number of samples for each class.

Weighted F1-score = (Support₁ × F1₁ + Support₂ × F1₂ + ... + Supportₙ × F1ₙ) / (Support₁ + Support₂ + ... + Supportₙ)

Where: Supportᵢ is the number of samples for class i, and F1ᵢ is the F1-score for class i.

SM 1.3. Downloading street-level images

We downloaded GSV images from public roads across the 23 study cities. We used OpenStreetMap data processed with the 'OSMnx' Python package v1.9.3 (Boeing, 2017) to identify all available public road lines in each city. We converted road lines from OSMnx into a set of points by sampling every 25 meters. We then queried GSV images adjacent to these points using the Google Maps Platform Street View Static API, in combination with the 'streetview' Python package (Letchford, 2024). We retained only images captured after 2016 with coordinates (i.e., latitude and longitude) and bearing (i.e., vehicle travel direction) information.

SM 1.4. Classifying the modal capture season into growing vs. dormant season

We identified street-level images within a 20-meter radius of each inventory tree's geographic coordinates. Then, we selected the most frequently occurring image capture month as the representative ("modal") month for that tree. To classify modal capture months into growing and dormant seasons, we obtained daily minimum and maximum air temperature data (1km resolution; Daymet v4) from 2016 to 2024 (Thornton et al., 2021). We picked this time frame because it corresponds to the temporal extent of the downloaded street-level imagery. Then, we extracted temperature values at each tree's coordinates using the 'terra' package (Hijmans, 2025).

Growing degree days (GDD), which are defined as the number of degrees where the average daily temperature exceeds a base threshold, were calculated as: GDD = Tmax+Tmin - Tbase (McMaster and Wilhelm, 1997). We used the Expert Team on Climate Change Detection and Indices of growing season length (ETCCDI), where the start of the season (SOS) was the first day with at least six consecutive days of GDD > 0. The end of the season (EOS) was the first day after July 1st with at least six consecutive days of GDD = 0. Median SOS and EOS were then computed across years per location and rounded from daily (MM-DD) to monthly (MM) resolution to match the temporal resolution of the downloaded street-level imagery. Lastly, we classified each tree's modal capture month as "growing" if it fell between median SOS and EOS, and "dormant" otherwise.

SM 1.5. Testing for spatial autocorrelation in tree-level matching among cities.

We anticipated spatial autocorrelation in the residuals of our generalized linear mixed models (GLMMs) because tree genera often exhibit clustered distributions influenced by local planting and environmental factors (McCoy et al., 2022), which can result in clustered residuals. Failing to account for this spatial structure may result in biased estimates and inflated Type I error rates (Legendre, 1993).

To test for and adequately address this autocorrelation, we calculated Global Moran’s *I* statistics on the GLMM residuals at distance intervals of 100 m up to 1,500 m (in 100 m increments). Given that we conducted hypothesis testing across multiple distance lags, we adjusted the p-values using a false discovery rate method to control for inflated Type I errors.

Initially, the spatial correlogram indicated weak but significant clustering at distances less than 1200 m, with no evidence of spatial structure beyond this distance. Subsequently, we fitted our GLMM using a random intercept based on our sampling grid IDs nested within city, extracted the residuals, and computed a spatial correlogram to account for the observed clustering (Figure S13). The results of our correlogram analysis indicated that no significant clustering remained in our residuals once the nested sampling grid ID random effect was included.

**Supplementary Figures**

Figure S1. Examples of AutoArborist images, iNaturalist images, and filtered iNaturalist images with Contrastive Language-Image Pre-Training (CLIP) deep learning model. Selected genera include *Acer* (maples), *Pinus* (pines), *Phoenix* (date palms), *Malus* (apples), and *Quercus* (oaks), which illustrate various tree growth forms. The left three columns contain examples of images for each genus from Google Street View (AutoArborist). The center three columns include examples of images for each genus from iNaturalist. The right three columns demonstrate the result of filtering iNaturalist images using the CLIP model with the text prompt “a photo of a large mature tree in an urban landscape”. These images were filtered to obtain a better correspondence between features visible in iNaturalist images and AutoArborist images and were subsequently used for training the tree classification model.

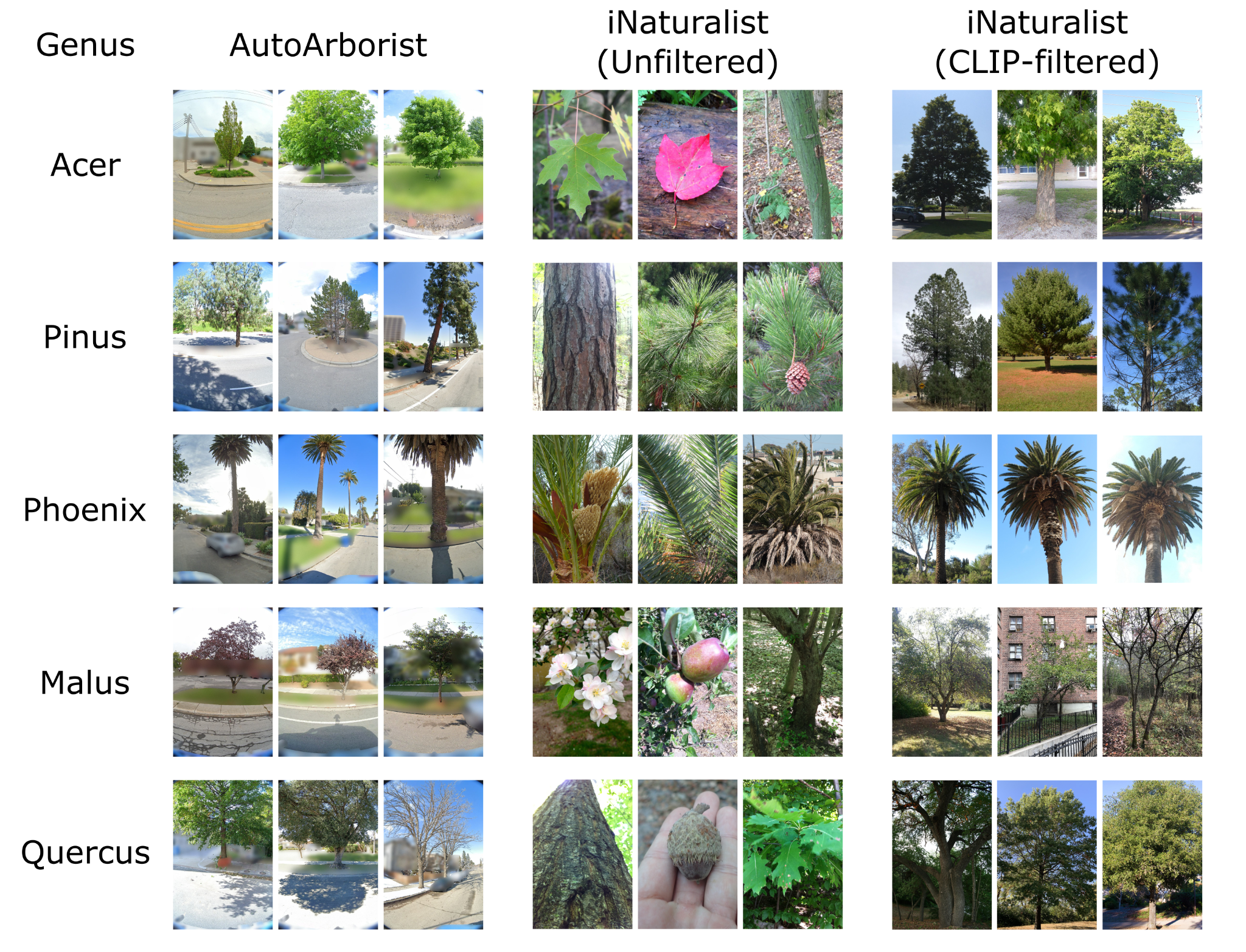

Figure S2. Location of Google Street View imagery included in the AutoArborist dataset for 23 study cities. Approximately 70% of images were allocated to model development (pink points), and the remaining 30% were withheld as a geographically independent test set to evaluate model generalizability (green points).

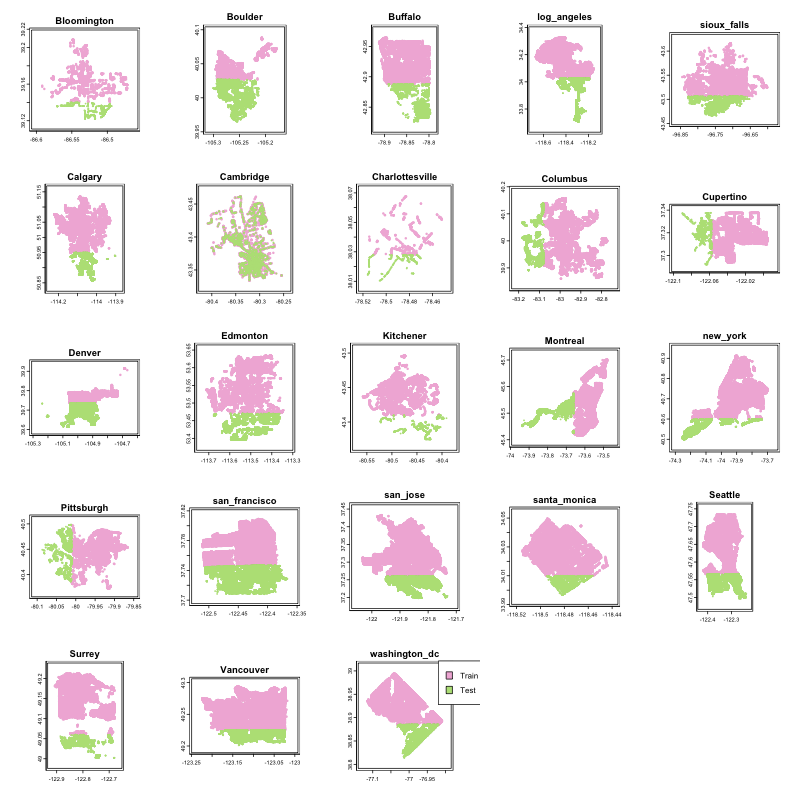

Figure S3. Diagnostic plots for regression models on genus Shannon diversity and classification performance, demonstrating the city of Kitchener, ON as an outlier.

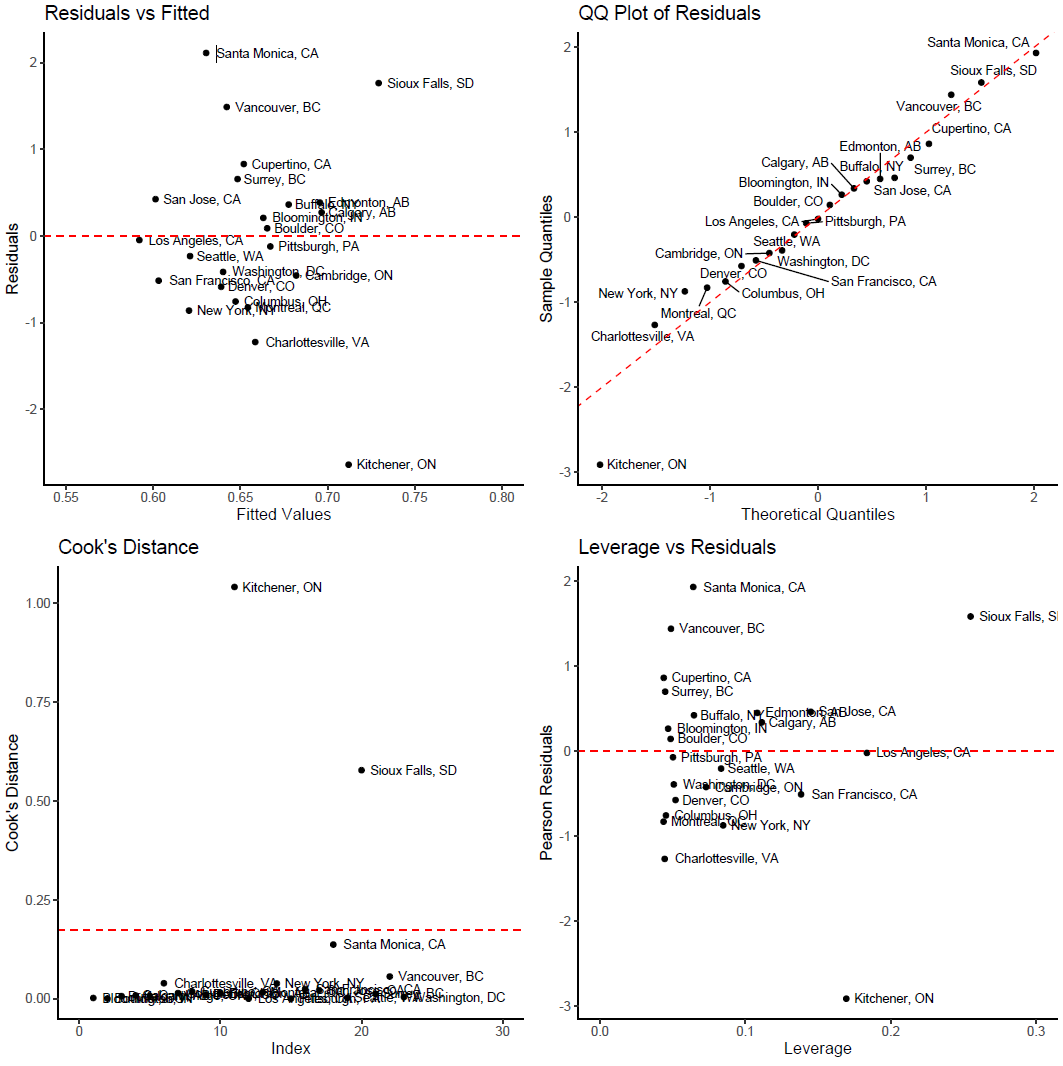

Figure S4. Diagnostic plots for regression models on genus richness and classification performance demonstrating the city of Kitchener, ON, as an outlier.

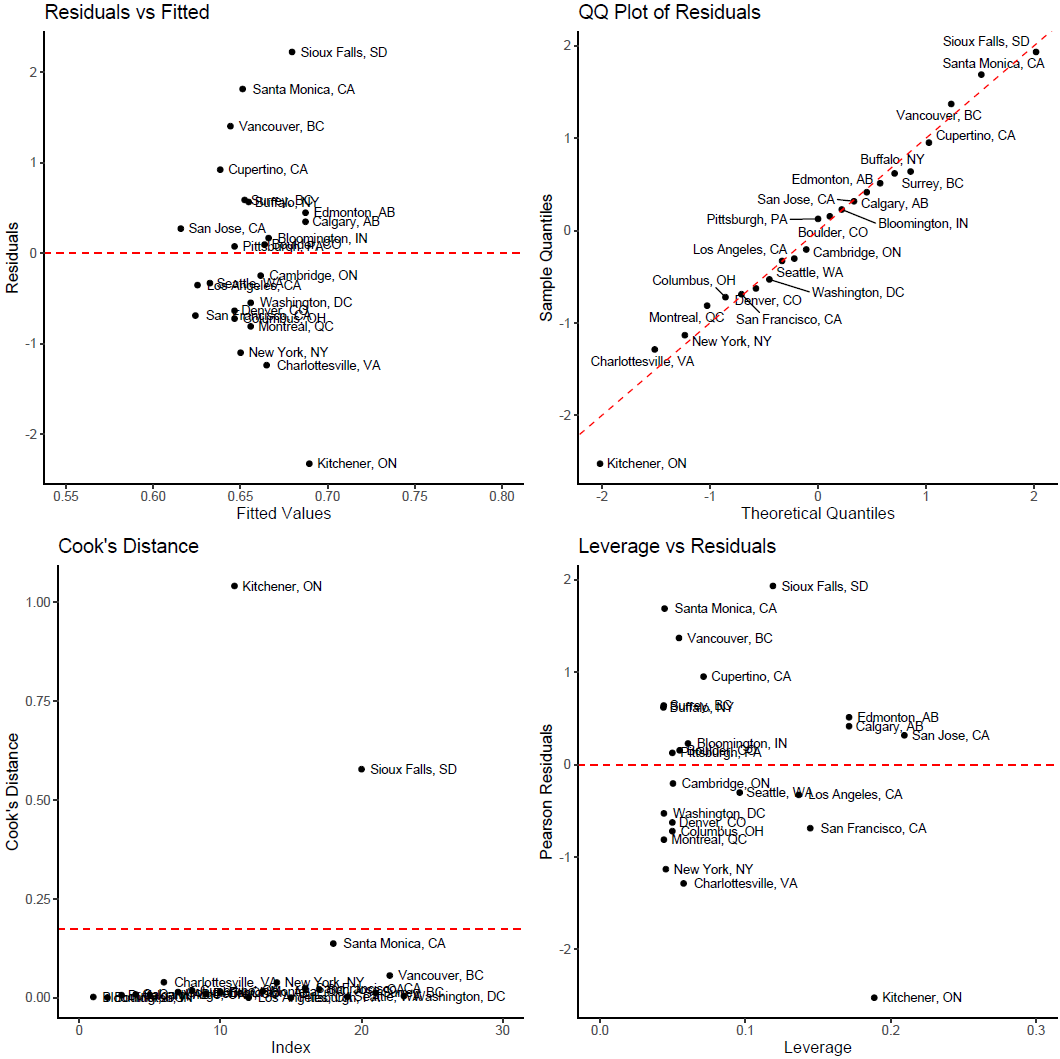

Figure S5. Diagnostic plots for regression models on genus evenness and classification performance demonstrating the city of Kitchener, ON, as an outlier.

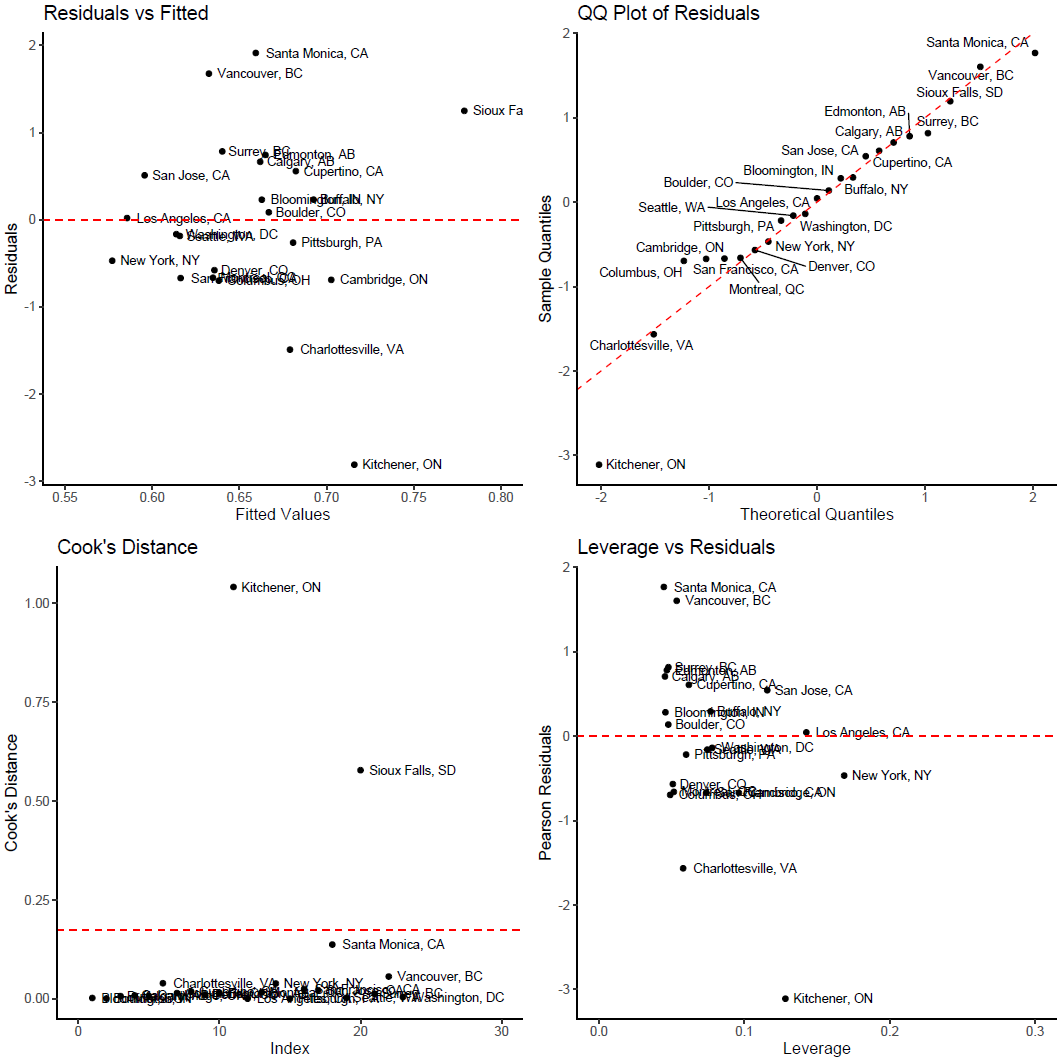

Figure S6. Map of 23 study cities and PCA biplot using WorldClim climate variables, grouping cities into four geographic regions.

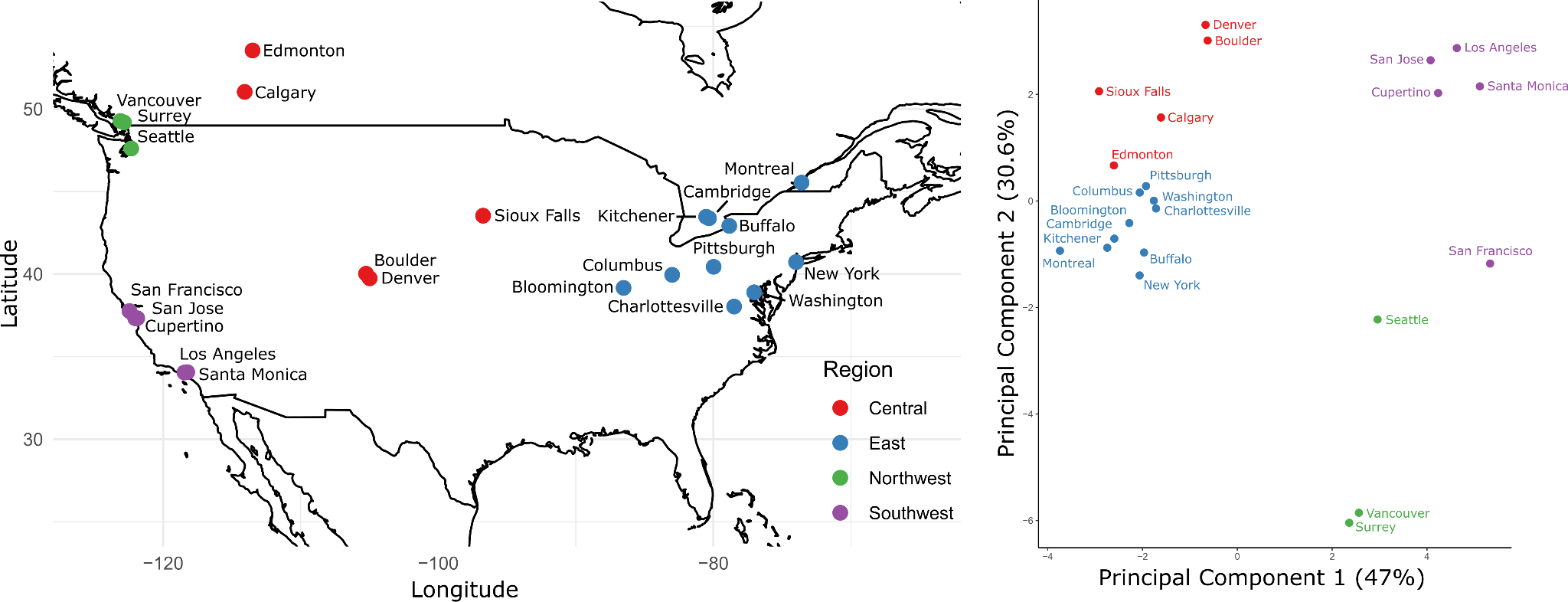

Figure S7. Precision and recall performance curve for the YOLOv5x tree detection model evaluated at the Intersection over Union (IOU) threshold of 0.5.

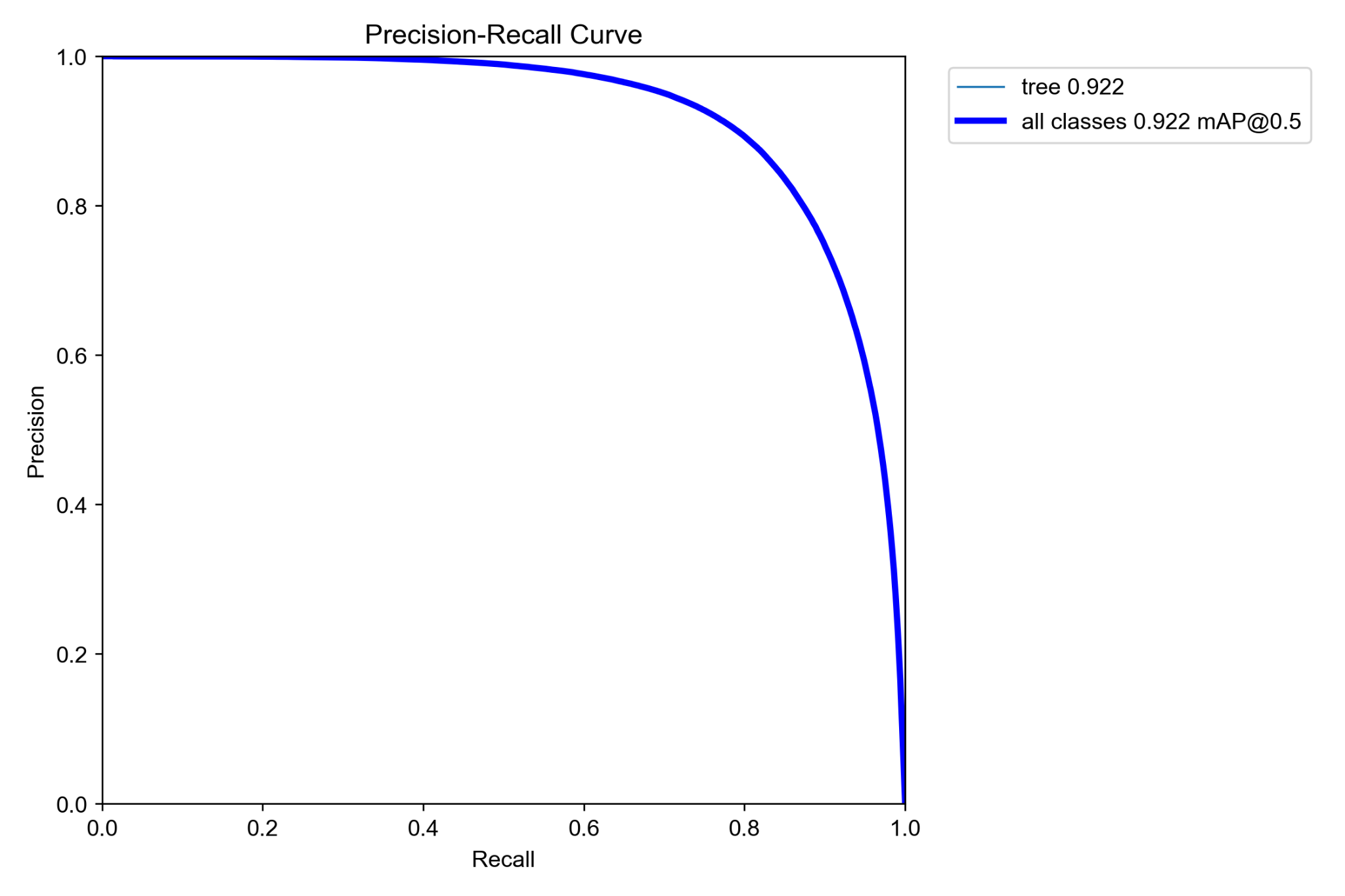

Figure S8. Scatterplot demonstrating the correlation between the number of training images used in the genus classification model and testing performance, as measured by F1 score. Points labelled by genus.

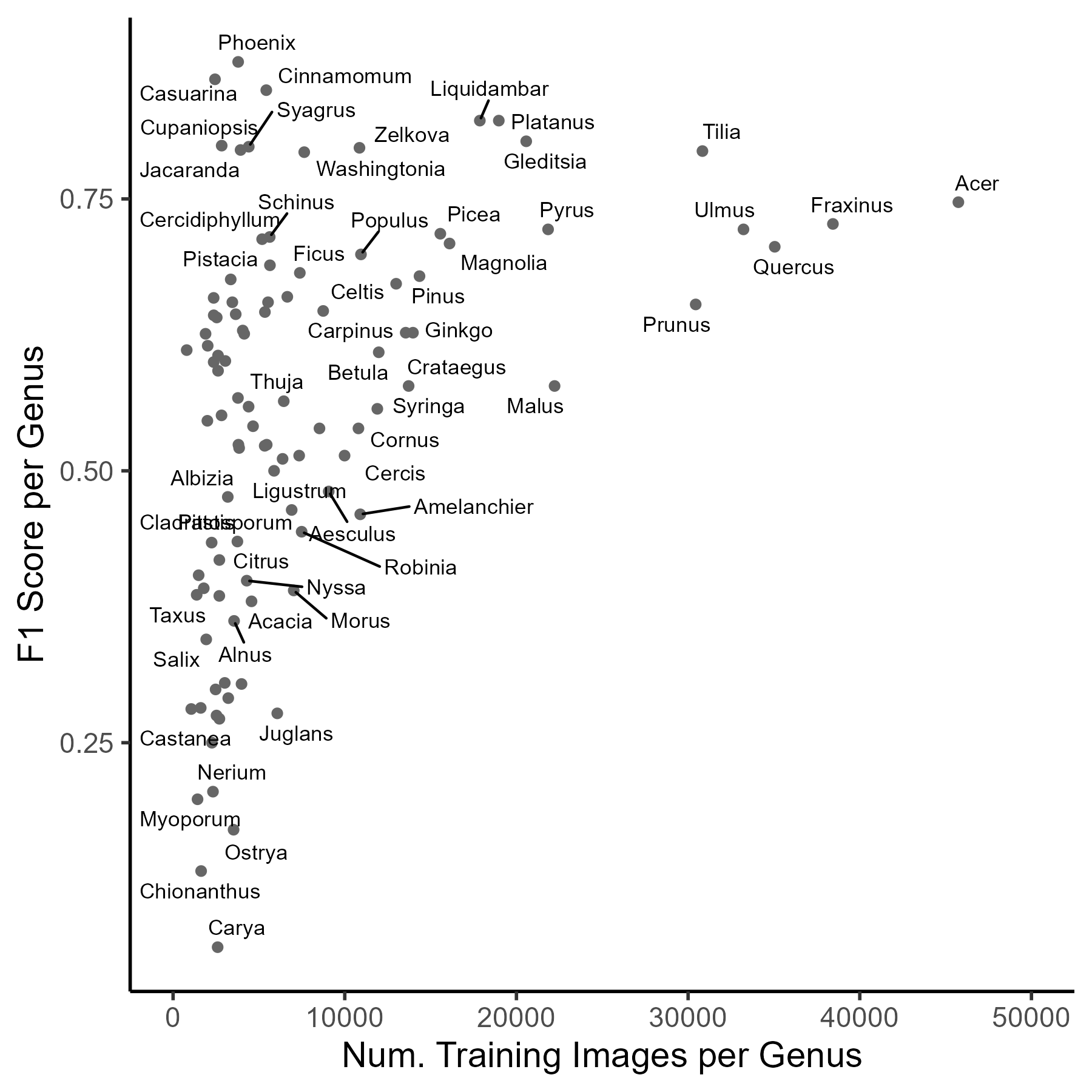

Figure S9. Effects of genus richness and evenness on classification performance across cities.

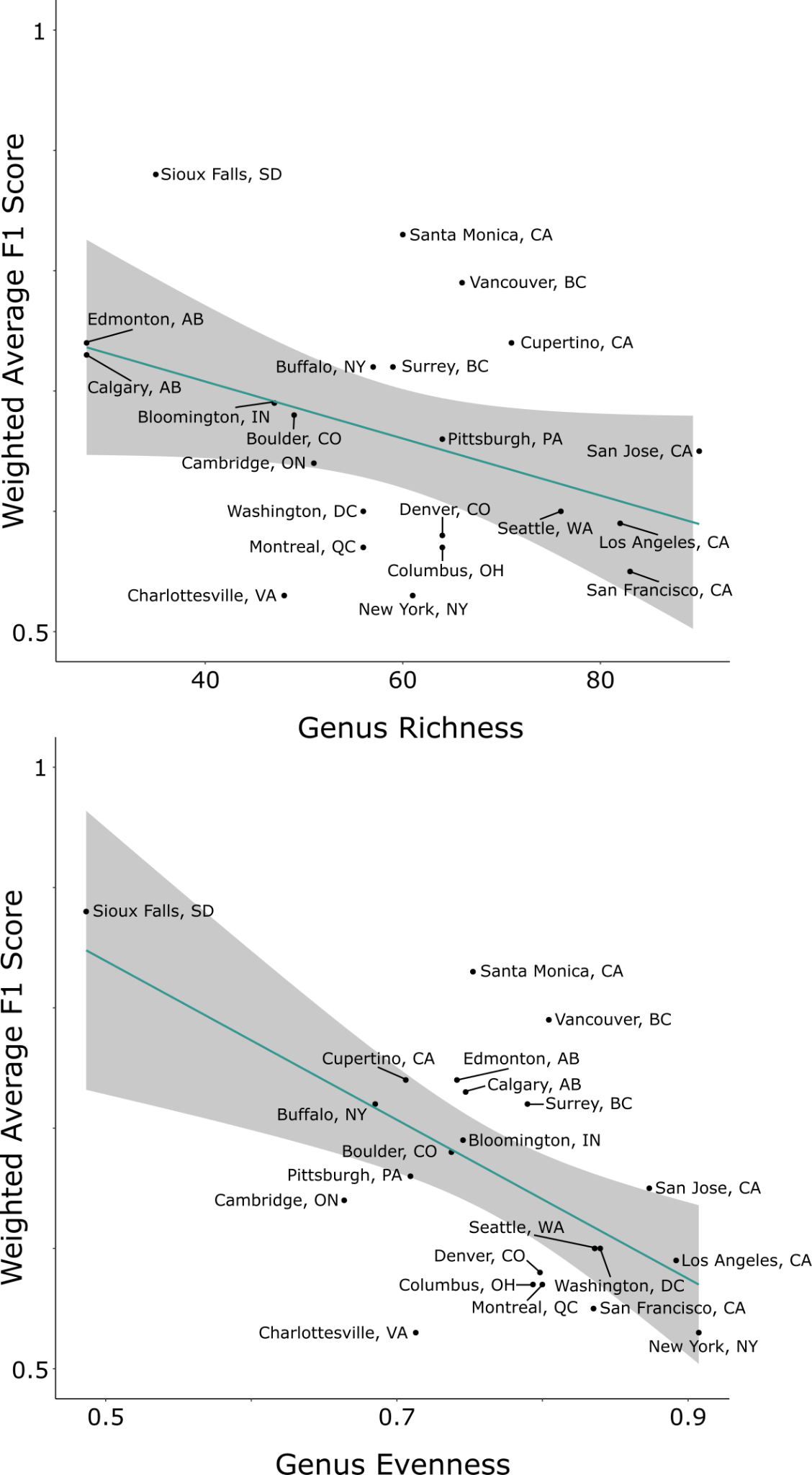

Figure S10. Differences in genus classification performance among regions.

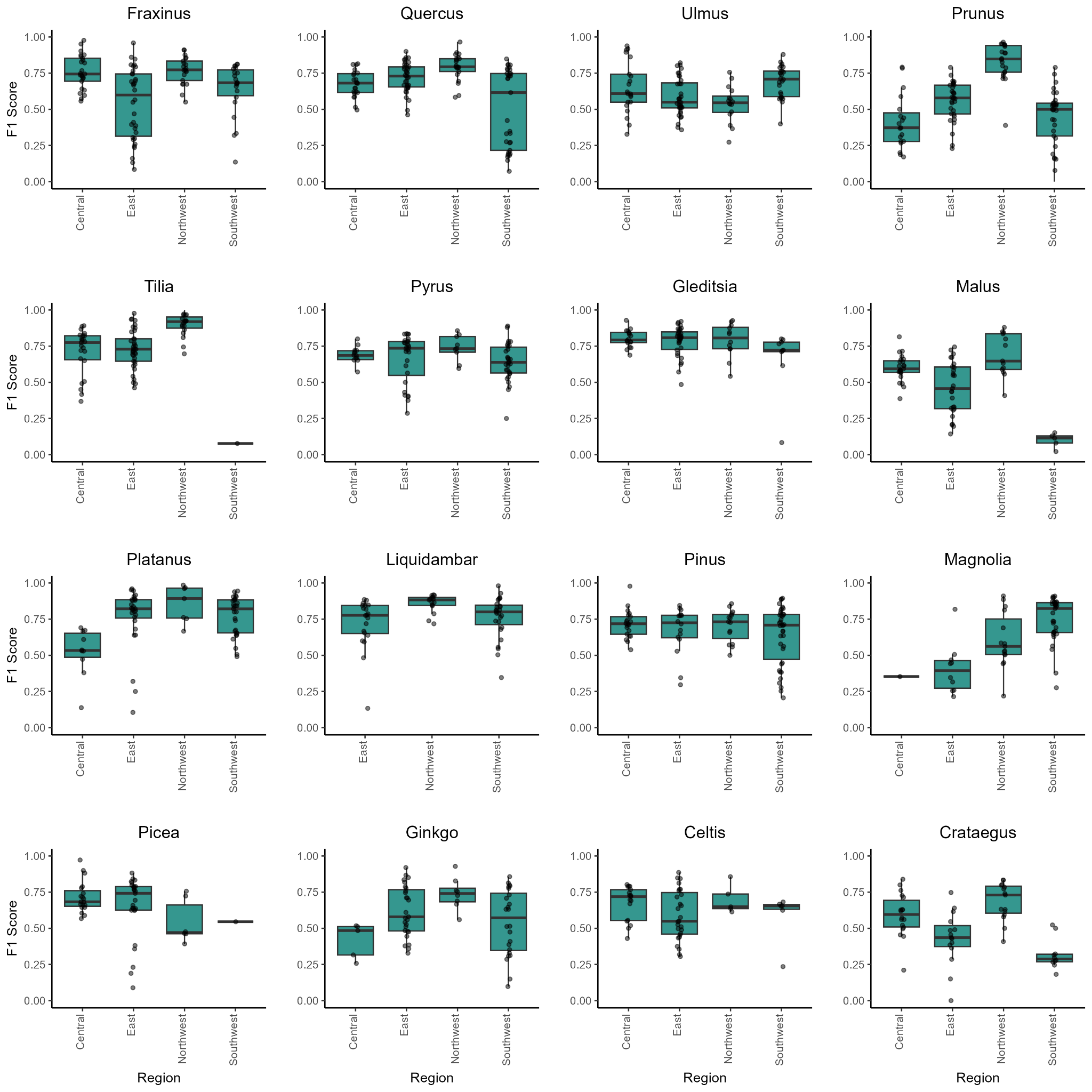

Figure S11. Differences in genus classification performance among months.

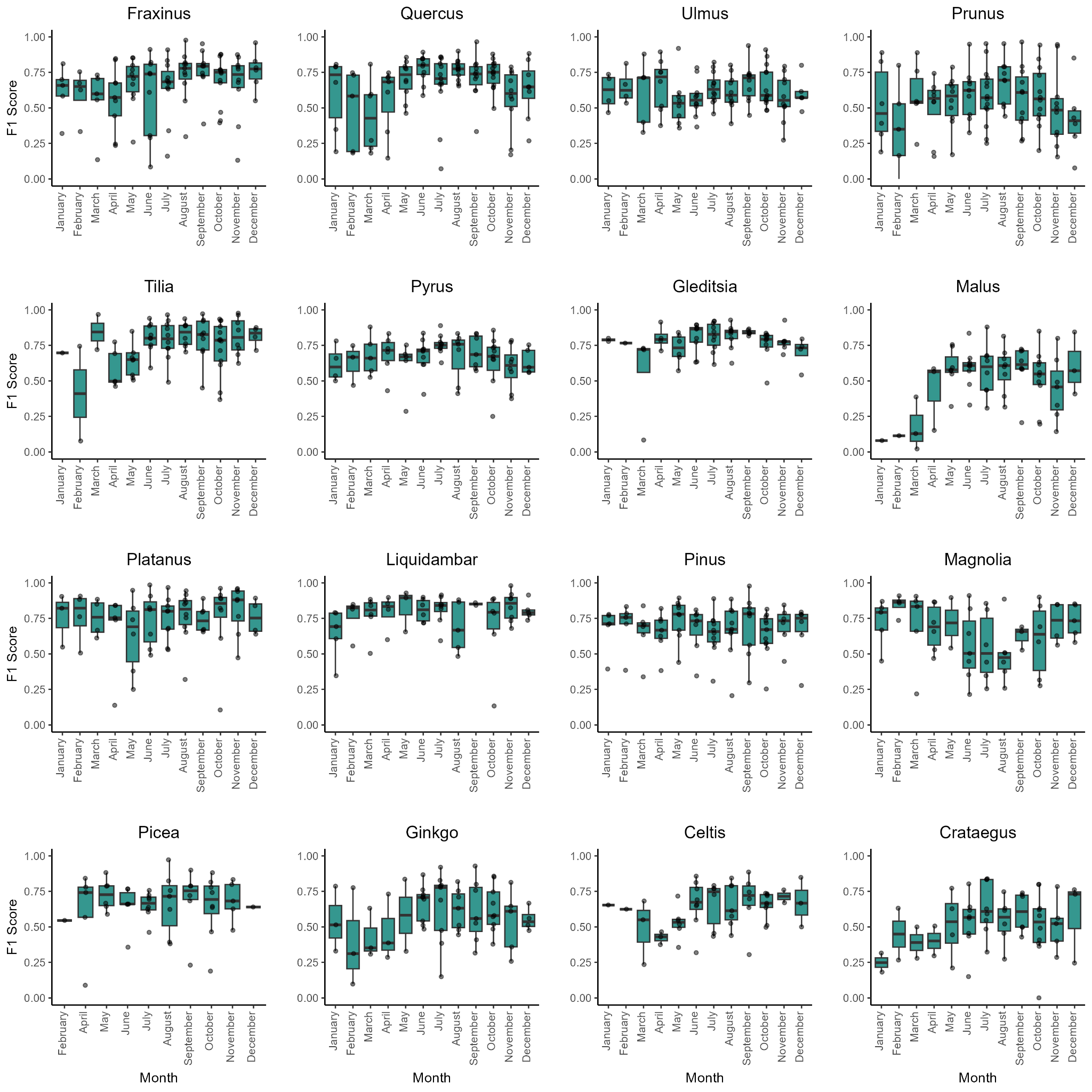

Figure S12. The proportion of inventory trees matched with at least one street-level image within the search radius, tested across radii of 5 m to 30 m, is shown overall and by city.

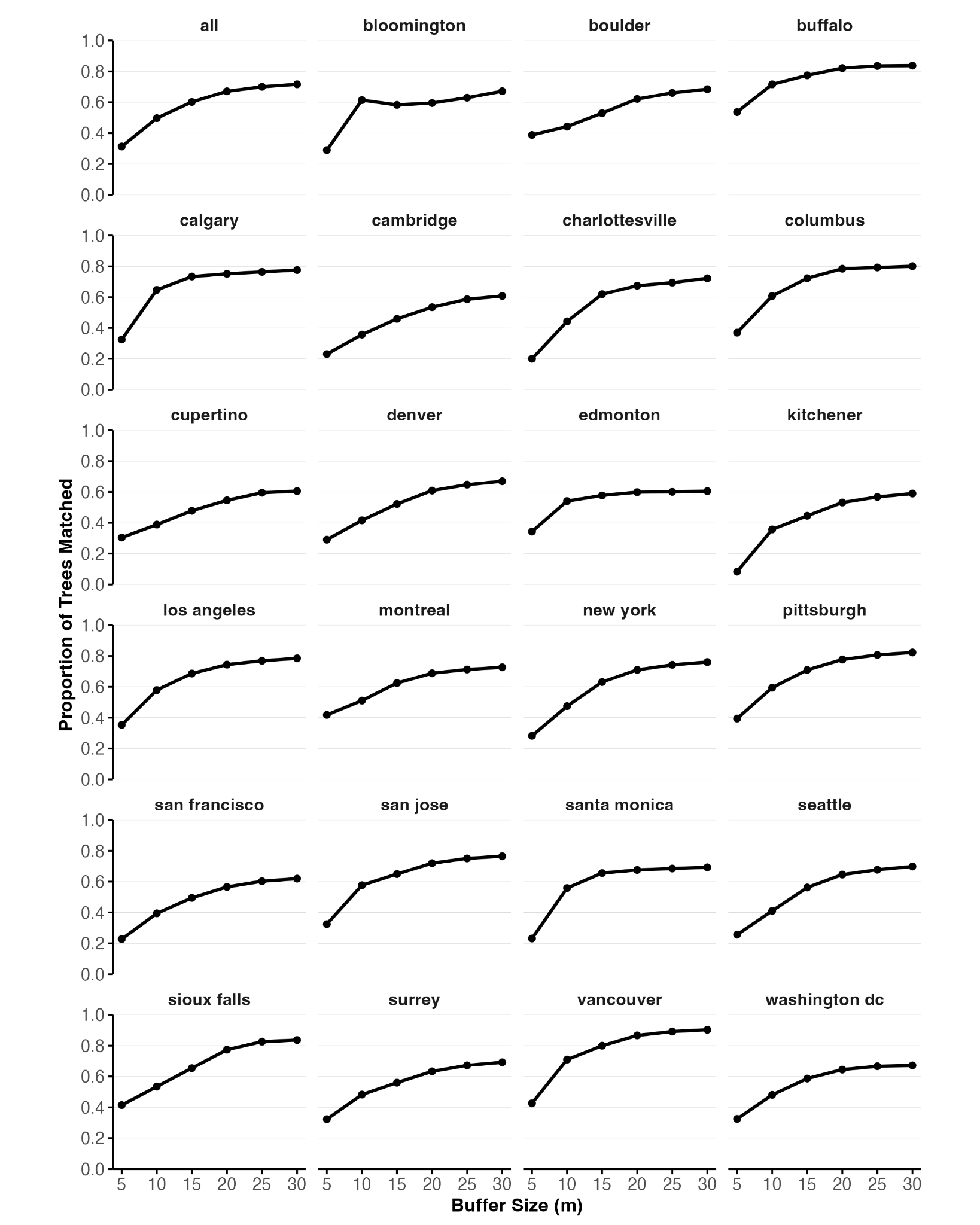

Figure S13. Residual diagnostics for the top binomial generalized linear mixed model (GLMM). (A) Quantile-quantile plot with the results of the uniformity (two-sample Kolmogorov-Smirnov) and outlier tests overlaid to assess distributional fit. (B) Simulated residuals (solid red line) versus predicted values with quantile regression fit (red-dotted line) and dispersion test results to evaluate heterogeneity and non-linearity.

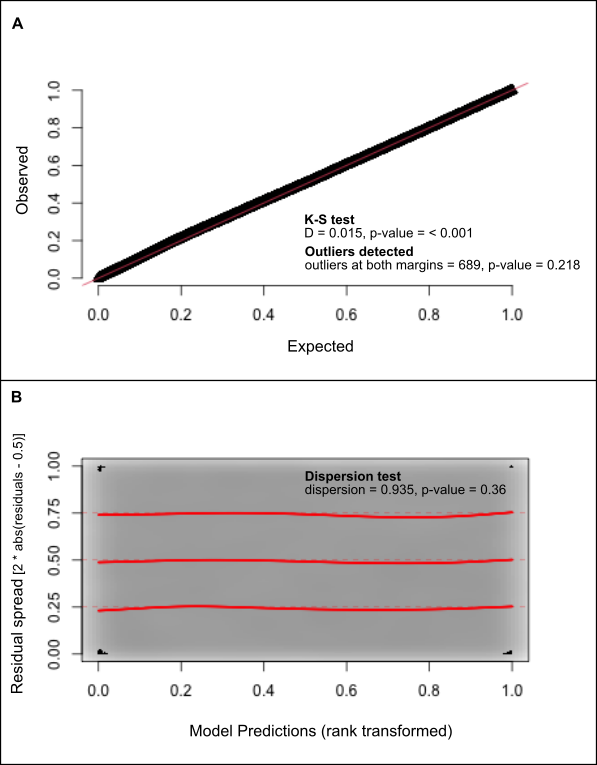

Figure S14. Correlogram of global Moran’s *I* for residuals from the top generalized linear mixed model (GLMM) across lag distances of 100-1500m (100m intervals) with and without the nested grid random intercept. Statistical significance of global Moran’s *I* was assessed via Monte Carlo permutation test (499 random permutations) with *p*-values corrected for multiple testing using the false discovery rate (FDR) method.

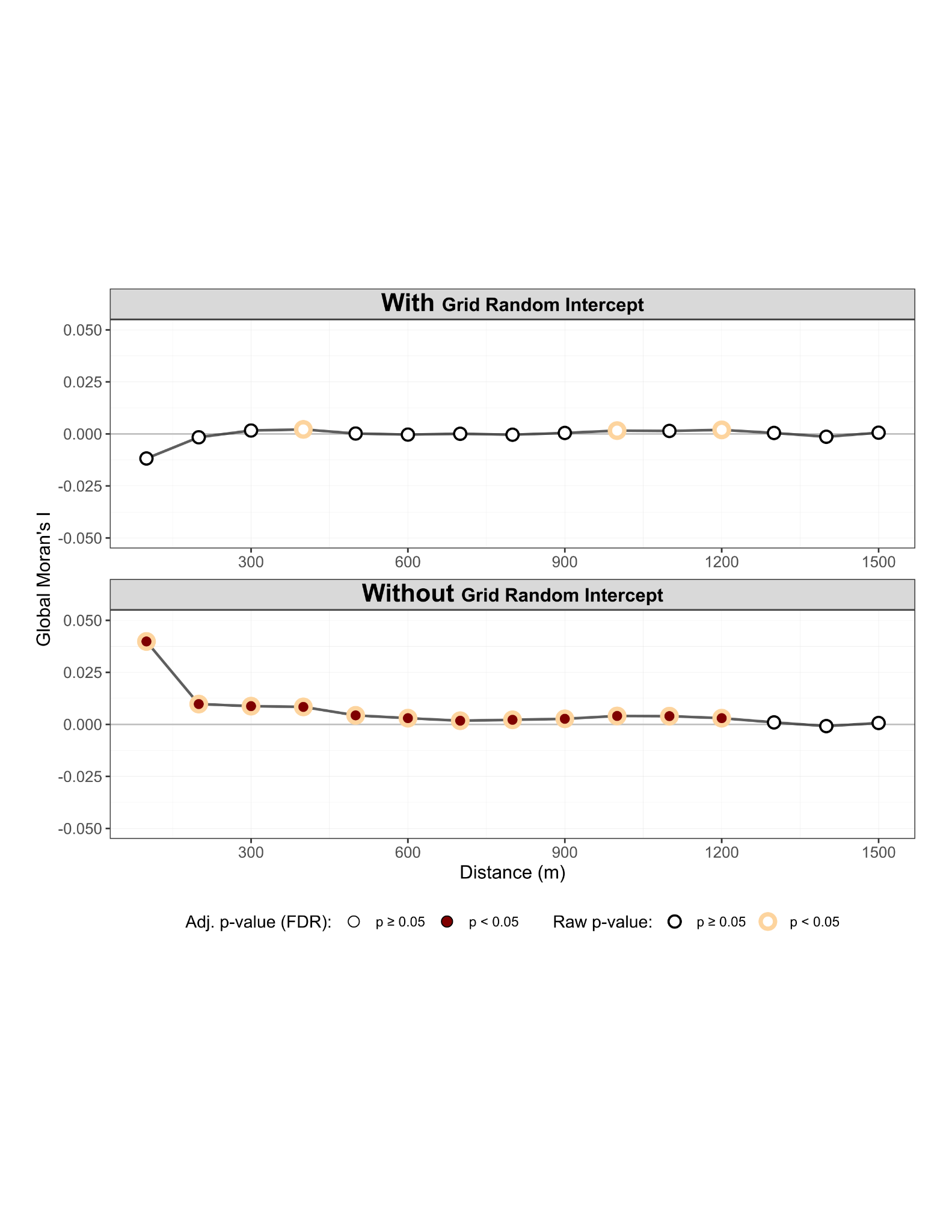

Figure S15. Best linear unbiased predictions (BLUPs) of random intercepts from the top generalized linear mixed model (GLMM). A) Genus-level BLUPs, showing variation in baseline tree matching probabilities across tree genera. B) City-level BLUPs, showing variation in baseline probabilities across cities. Points represent log-odds estimates, and 95% confidence intervals are shown as error bars. Significant deviations (over/under-performance) from zero are colored with filled point estimates (orange for city, teal for genus). In contrast, non-significant deviations are shown in grey with hollow point estimates.

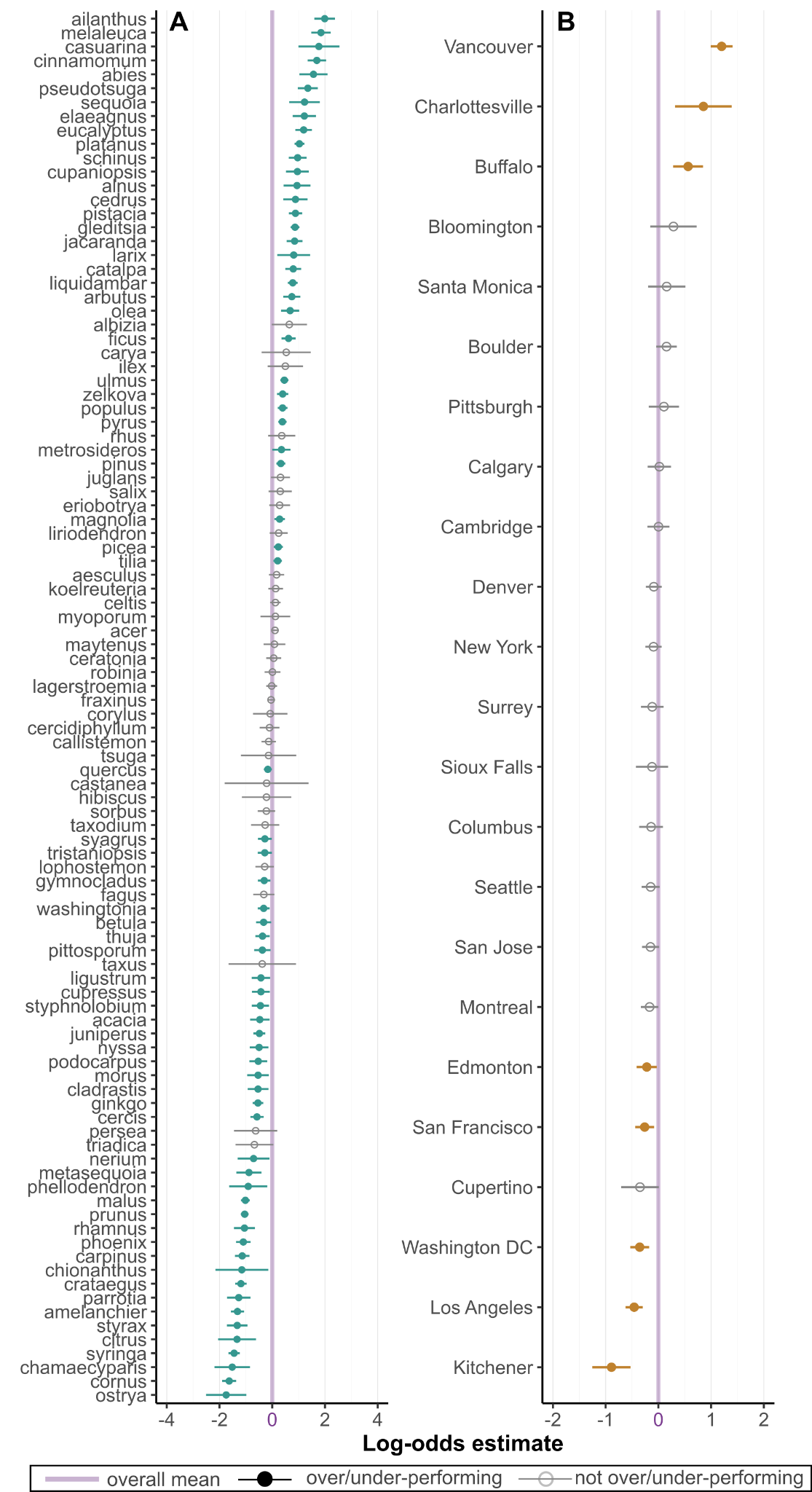

**Supplementary Tables**

Table S1. Selected 100 tree genera examined in the study, and the number of training, validation, and testing images and sources used in the tree classification model.

| Genus | Common Name | Training Images (AutoArborist) | Training Images (iNaturalist) | Total Training Images | Validation Images | Testing Images (10% Random Split) | Testing Images (30% Geographic Split) |
| --- | --- | --- | --- | --- | --- | --- | --- |
| Acer | Maple | 44743 | 1000 | 45743 | 4574 | 4934 | 33450 |
| Fraxinus | Ash | 37435 | 1000 | 38435 | 3843 | 4067 | 23652 |
| Quercus | Oak | 34047 | 1000 | 35047 | 3504 | 3534 | 19970 |
| Ulmus | Elm | 32226 | 1000 | 33226 | 3322 | 3457 | 20617 |
| Tilia | Basswood | 29835 | 1000 | 30835 | 3083 | 3278 | 14260 |
| Prunus | Cherry / Plum | 29442 | 1000 | 30442 | 3044 | 3231 | 21906 |
| Malus | Apple | 21943 | 279 | 22222 | 2222 | 2439 | 9339 |
| Pyrus | Pear | 20844 | 1000 | 21844 | 2184 | 2274 | 12798 |
| Gleditsia | Honeylocust | 19570 | 1000 | 20570 | 2057 | 2009 | 12885 |
| Platanus | Sycamore | 18450 | 523 | 18973 | 1897 | 2055 | 10041 |
| Liquidambar | Sweetgum | 16868 | 1000 | 17868 | 1786 | 1775 | 8676 |
| Magnolia | Magnolia | 15104 | 1000 | 16104 | 1610 | 1555 | 7368 |
| Picea | Spurce | 14567 | 1000 | 15567 | 1556 | 1618 | 6708 |
| Pinus | Pine | 14077 | 277 | 14354 | 1435 | 1496 | 9171 |
| Ginkgo | Ginkgo | 13946 | 26 | 13972 | 1397 | 1548 | 5146 |
| Crataegus | Hawthorn | 12720 | 1000 | 13720 | 1372 | 1283 | 4514 |
| Carpinus | Hornbeam | 12550 | 1000 | 13550 | 1355 | 1256 | 3146 |
| Celtis | Hackberry | 12142 | 850 | 12992 | 1299 | 1217 | 5462 |
| Syringa | Lilac | 11281 | 616 | 11897 | 1189 | 1154 | 4068 |
| Betula | Birch | 10979 | 1000 | 11979 | 1197 | 1078 | 2104 |
| Zelkova | Zelkova | 10836 | 21 | 10857 | 1085 | 1218 | 6019 |
| Populus | Poplar / Aspen | 9944 | 1000 | 10944 | 1094 | 997 | 6527 |
| Amelanchier | Serviceberry | 9903 | 1000 | 10903 | 1090 | 1023 | 2959 |
| Cornus | Dogwood | 9797 | 1000 | 10797 | 1079 | 983 | 2904 |
| Cercis | Redbug | 8989 | 1000 | 9989 | 998 | 894 | 2710 |
| Lagerstroemia | Crape Myrtle | 8701 | 43 | 8744 | 874 | 929 | 3815 |
| Aesculus | Horse Chestnut | 8065 | 1000 | 9065 | 906 | 759 | 1581 |
| Gymnocladus | Kentucky Coffeetree | 8059 | 468 | 8527 | 852 | 827 | 2710 |
| Washingtonia | Fan Palm | 6953 | 686 | 7639 | 763 | 729 | 3674 |
| Robinia | Black Locust | 6489 | 1000 | 7489 | 748 | 629 | 1614 |
| Ficus | Fig | 6383 | 1000 | 7383 | 738 | 583 | 3170 |
| Liriodendron | Tulip Tree | 6344 | 1000 | 7344 | 734 | 644 | 1175 |
| Koelreuteria | Goldenrain Tree | 6108 | 271 | 6379 | 637 | 679 | 1694 |
| Morus | Mulberry | 6027 | 1000 | 7027 | 702 | 553 | 816 |
| Ligustrum | Privet | 5908 | 1000 | 6908 | 690 | 552 | 1542 |
| Fagus | Beech | 5653 | 1000 | 6653 | 665 | 523 | 1282 |
| Thuja | Arborvitae | 5446 | 1000 | 6446 | 644 | 520 | 1958 |
| Pistacia | Pistache | 5394 | 247 | 5641 | 564 | 572 | 2967 |
| Catalpa | Catalpa | 5269 | 262 | 5531 | 553 | 558 | 1503 |
| Cinnamomum | Camphor Tree | 5244 | 184 | 5428 | 542 | 526 | 3075 |
| Cercidiphyllum | Katsura Tree | 5162 | 25 | 5187 | 518 | 531 | 1046 |
| Eucalyptus | Eucalpytus | 5071 | 278 | 5349 | 534 | 565 | 3548 |
| Juglans | Walnut | 5069 | 1000 | 6069 | 606 | 441 | 463 |
| Sorbus | Mountain Ash | 4882 | 1000 | 5882 | 588 | 407 | 1068 |
| Juniperus | Juniper | 4775 | 561 | 5336 | 533 | 434 | 3296 |
| Cupressus | Cypress | 4656 | 1 | 4657 | 465 | 505 | 2210 |
| Schinus | Pepper Tree | 4628 | 1000 | 5628 | 562 | 392 | 2337 |
| Ailanthus | Tree of Heaven | 4453 | 1000 | 5453 | 545 | 296 | 974 |
| Syagrus | Queen Palm | 4392 | 16 | 4408 | 440 | 534 | 3114 |
| Parrotia | Persian Ironwood | 4143 | 0 | 4143 | 414 | 427 | 979 |
| Jacaranda | Jacaranda | 3927 | 4 | 3931 | 393 | 431 | 2457 |
| Nyssa | Tupelo | 3790 | 501 | 4291 | 429 | 342 | 1234 |
| Olea | Olive | 3699 | 148 | 3847 | 384 | 378 | 1425 |
| Acacia | Acacia | 3681 | 888 | 4569 | 456 | 305 | 1601 |
| Cedrus | Cedar | 3641 | 7 | 3648 | 364 | 384 | 596 |
| Phoenix | Date Palm | 3614 | 176 | 3790 | 379 | 391 | 2386 |
| Pittosporum | Pittosporum | 3585 | 161 | 3746 | 374 | 392 | 2264 |
| Styrax | Snowbell | 3501 | 305 | 3806 | 380 | 350 | 782 |
| Ceratonia | Carob | 3408 | 43 | 3451 | 345 | 340 | 2877 |
| Arbutus | Madrones | 3399 | 1000 | 4399 | 439 | 272 | 2195 |
| Callistemon | Bottlebrush | 3357 | 0 | 3357 | 335 | 340 | 3520 |
| Sequoia | Coast Redrood | 3059 | 1000 | 4059 | 405 | 244 | 474 |
| Taxodium | Bald Cypress | 2986 | 1000 | 3986 | 398 | 201 | 349 |
| Melaleuca | Paperbark | 2862 | 181 | 3043 | 304 | 316 | 2021 |
| Pseudotsuga | Douglas Fir | 2778 | 1000 | 3778 | 377 | 210 | 766 |
| Cupaniopsis | Tuckeroo | 2707 | 124 | 2831 | 283 | 286 | 2183 |
| Eriobotrya | Loquat | 2688 | 134 | 2822 | 282 | 287 | 963 |
| Tristaniopsis | Water Bum | 2618 | 0 | 2618 | 261 | 288 | 2339 |
| Metasequoia | Dawn Redwood | 2601 | 18 | 2619 | 261 | 310 | 631 |
| Alnus | Alder | 2553 | 1000 | 3553 | 355 | 180 | 527 |
| Ostrya | Hop-hornbeam | 2524 | 1000 | 3524 | 352 | 182 | 352 |
| Podocarpus | Podocarpus | 2351 | 12 | 2363 | 236 | 282 | 1877 |
| Casuarina | She-oak | 2317 | 124 | 2441 | 244 | 240 | 394 |
| Citrus | Citrus | 2228 | 467 | 2695 | 269 | 205 | 333 |
| Corylus | Hazel | 2213 | 1000 | 3213 | 321 | 133 | 333 |
| Albizia | Silk Tree | 2186 | 1000 | 3186 | 318 | 133 | 258 |
| Cladrastis | Yellowwood | 2170 | 76 | 2246 | 224 | 235 | 1110 |
| Nerium | Oleander | 2066 | 257 | 2323 | 232 | 208 | 260 |
| Ilex | Holly | 2010 | 1000 | 3010 | 301 | 97 | 223 |
| Lophostemon | Brush Box | 1996 | 1 | 1997 | 199 | 222 | 1807 |
| Styphnolobium | Japanese Pagoda Tree | 1959 | 405 | 2364 | 236 | 177 | 1218 |
| Maytenus | Mayten | 1934 | 82 | 2016 | 201 | 176 | 759 |
| Metrosideros | New Zealand Christmas Tree | 1886 | 5 | 1891 | 189 | 219 | 2203 |
| Triadica | Chinese Tallow Tree | 1855 | 513 | 2368 | 236 | 160 | 347 |
| Rhamnus | Buckthorn | 1697 | 1000 | 2697 | 269 | 79 | 1065 |
| Rhus | Sumac | 1692 | 1000 | 2692 | 269 | 62 | 565 |
| Carya | Hickory | 1595 | 1000 | 2595 | 259 | 59 | 132 |
| Elaeagnus | Silverberry | 1551 | 992 | 2543 | 254 | 80 | 556 |
| Tsuga | Hemlock | 1530 | 1000 | 2530 | 253 | 45 | 89 |
| Salix | Willow | 1518 | 412 | 1930 | 193 | 106 | 509 |
| Abies | Fir | 1476 | 1000 | 2476 | 247 | 52 | 273 |
| Chamaecyparis | False Cypress | 1471 | 18 | 1489 | 148 | 171 | 384 |
| Phellodendron | Amur Cork Tree | 1401 | 209 | 1610 | 161 | 124 | 226 |
| Castanea | Chestnut | 1262 | 1000 | 2262 | 226 | 28 | 25 |
| Persea | Avocado | 1253 | 534 | 1787 | 178 | 84 | 202 |
| Myoporum | Myoporum | 1221 | 203 | 1424 | 142 | 121 | 456 |
| Chionanthus | Fringe Tree | 1062 | 571 | 1633 | 163 | 45 | 183 |
| Taxus | Yew | 811 | 563 | 1374 | 137 | 31 | 174 |
| Hibiscus | Hibiscus | 787 | 271 | 1058 | 105 | 57 | 194 |
| Larix | Larch | 700 | 93 | 793 | 79 | 65 | 233 |
| Total | - | 730688 | 59132 | 789820 | 78934 | 74613 | 356341 |

Table S2. Counts of grids, genera, and AutoArborist inventory trees per city and overall. The “without buffer” dataset includes all stratified sampled inventory trees to which the computer-vision-based tree inventory approach was applied, whereas the “with buffer” dataset is the subset where at least one Google Street View image was located within the 20-m search buffer. The “with buffer” dataset was used in the binomial generalized linear mixed model (GLMM) analysis.

| **City** | **without buffer** | | |  | **with buffer** | | |
| --- | --- | --- | --- | --- | --- | --- | --- |
|  | **N Grid** | **N Genus** | **N Trees** |  | **N Grid** | **N Genus** | **N Trees** |
| Bloomington | 21 | 29 | 332 |  | 21 | 29 | 331 |
| Boulder | 193 | 46 | 3790 |  | 193 | 44 | 3523 |
| Buffalo | 102 | 39 | 1112 |  | 100 | 36 | 936 |
| Calgary | 148 | 19 | 2792 |  | 147 | 18 | 2474 |
| Cambridge | 147 | 44 | 3363 |  | 140 | 43 | 2857 |
| Charlottesville | 12 | 26 | 151 |  | 11 | 24 | 132 |
| Columbus | 281 | 45 | 2585 |  | 184 | 39 | 1469 |
| Cupertino | 36 | 40 | 533 |  | 33 | 38 | 458 |
| Denver | 745 | 62 | 17305 |  | 728 | 62 | 15721 |
| Edmonton | 211 | 22 | 5735 |  | 206 | 20 | 4657 |
| Kitchener | 30 | 17 | 434 |  | 30 | 17 | 416 |
| Los Angeles | 1046 | 64 | 11479 |  | 805 | 64 | 8519 |
| Montreal | 437 | 49 | 7754 |  | 422 | 49 | 6735 |
| New York | 624 | 58 | 13835 |  | 621 | 58 | 13219 |
| Pittsburgh | 76 | 36 | 1038 |  | 74 | 34 | 986 |
| San Francisco | 367 | 73 | 5427 |  | 363 | 72 | 4908 |
| San Jose | 605 | 75 | 8566 |  | 582 | 75 | 7757 |
| Santa Monica | 42 | 34 | 722 |  | 42 | 33 | 552 |
| Seattle | 402 | 70 | 5335 |  | 401 | 70 | 5207 |
| Sioux Falls | 85 | 20 | 633 |  | 85 | 17 | 544 |
| Surrey | 161 | 56 | 2382 |  | 160 | 45 | 1998 |
| Vancouver | 233 | 49 | 3046 |  | 229 | 48 | 2923 |
| Washington DC | 302 | 49 | 5079 |  | 287 | 47 | 4367 |
| **Totals:** | **6306** |  | **103428** |  | **5864** |  | **90689** |

Table S3. Performance metrics for image classification model of 100 tree genera sorted by support (the number of testing images).

| Genus | Common Name | Precision | Recall | F1-Score | Support |
| --- | --- | --- | --- | --- | --- |
| Acer | Maple | 0.791 | 0.708 | 0.747 | 33450 |
| Fraxinus | Ash | 0.729 | 0.724 | 0.727 | 23652 |
| Prunus | Cherry / Plum | 0.692 | 0.618 | 0.653 | 21906 |
| Ulmus | Elm | 0.706 | 0.740 | 0.722 | 20617 |
| Quercus | Oak | 0.715 | 0.698 | 0.706 | 19970 |
| Tilia | Basswood | 0.798 | 0.790 | 0.794 | 14260 |
| Gleditsia | Honey Locust | 0.812 | 0.794 | 0.803 | 12885 |
| Pyrus | Pear | 0.715 | 0.728 | 0.722 | 12798 |
| Platanus | Sycamore | 0.844 | 0.800 | 0.822 | 10041 |
| Malus | Apple | 0.525 | 0.642 | 0.578 | 9339 |
| Pinus | Pine | 0.708 | 0.652 | 0.679 | 9171 |
| Liquidambar | Sweetgum | 0.858 | 0.789 | 0.822 | 8676 |
| Magnolia | Magnolia | 0.670 | 0.753 | 0.709 | 7368 |
| Picea | Spruce | 0.673 | 0.768 | 0.718 | 6708 |
| Populus | Poplar / Aspen | 0.641 | 0.768 | 0.699 | 6527 |
| Zelkova | Zelkova | 0.839 | 0.759 | 0.797 | 6019 |
| Celtis | Hackberry | 0.677 | 0.667 | 0.672 | 5462 |
| Ginkgo | Ginkgo | 0.557 | 0.718 | 0.627 | 5146 |
| Crataegus | Hawthorn | 0.576 | 0.579 | 0.578 | 4514 |
| Syringa | Lilac | 0.597 | 0.523 | 0.557 | 4068 |
| Lagerstroemia | Crape Myrtle | 0.617 | 0.680 | 0.647 | 3815 |
| Washingtonia | Fan Palm | 0.766 | 0.823 | 0.793 | 3674 |
| Eucalyptus | Eucalyptus | 0.694 | 0.605 | 0.646 | 3548 |
| Callistemon | Bottlebrush | 0.682 | 0.671 | 0.676 | 3520 |
| Juniperus | Juniper | 0.567 | 0.485 | 0.523 | 3296 |
| Ficus | Fig | 0.699 | 0.667 | 0.682 | 3170 |
| Carpinus | Hornbeam | 0.619 | 0.635 | 0.627 | 3146 |
| Syagrus | Queen Palm | 0.771 | 0.827 | 0.798 | 3114 |
| Cinnamomum | Camphor Tree | 0.845 | 0.856 | 0.850 | 3075 |
| Pistacia | Pistache | 0.663 | 0.718 | 0.689 | 2967 |
| Amelanchier | Serviceberry | 0.433 | 0.491 | 0.460 | 2959 |
| Cornus | Dogwood | 0.613 | 0.480 | 0.539 | 2904 |
| Ceratonia | Carob | 0.707 | 0.610 | 0.655 | 2877 |
| Cercis | Redbud | 0.504 | 0.524 | 0.514 | 2710 |
| Gymnocladus | Kentucky Coffeetree | 0.489 | 0.601 | 0.539 | 2710 |
| Jacaranda | Jacaranda | 0.845 | 0.752 | 0.795 | 2457 |
| Phoenix | Date Palm | 0.861 | 0.892 | 0.876 | 2386 |
| Tristaniopsis | Water Gum | 0.642 | 0.574 | 0.606 | 2339 |
| Schinus | Pepper Tree | 0.762 | 0.674 | 0.715 | 2337 |
| Pittosporum | Pittosporum | 0.436 | 0.435 | 0.435 | 2264 |
| Cupressus | Cypress | 0.480 | 0.620 | 0.541 | 2210 |
| Metrosideros | New Zealand Christmas Tree | 0.589 | 0.667 | 0.626 | 2203 |
| Arbutus | Madrone | 0.557 | 0.561 | 0.559 | 2195 |
| Cupaniopsis | Tuckeroo | 0.815 | 0.784 | 0.799 | 2183 |
| Betula | Birch | 0.582 | 0.639 | 0.609 | 2104 |
| Melaleuca | Paperbark | 0.562 | 0.646 | 0.601 | 2021 |
| Thuja | Arborvitae | 0.529 | 0.603 | 0.564 | 1958 |
| Podocarpus | Podocarpus | 0.752 | 0.587 | 0.659 | 1877 |
| Lophostemon | Brush Box | 0.568 | 0.527 | 0.546 | 1807 |
| Koelreuteria | Goldenrain Tree | 0.458 | 0.579 | 0.511 | 1694 |
| Robinia | Black Locust | 0.438 | 0.450 | 0.444 | 1614 |
| Acacia | Acacia | 0.368 | 0.394 | 0.380 | 1601 |
| Aesculus | Horse Chestnut | 0.502 | 0.462 | 0.481 | 1581 |
| Ligustrum | Privet | 0.419 | 0.519 | 0.464 | 1542 |
| Catalpa | Catalpa | 0.632 | 0.679 | 0.655 | 1503 |
| Olea | Olive | 0.471 | 0.584 | 0.521 | 1425 |
| Fagus | Beech | 0.636 | 0.686 | 0.660 | 1282 |
| Nyssa | Tupelo | 0.362 | 0.443 | 0.399 | 1234 |
| Styphnolobium | Japanese Pagoda Tree | 0.711 | 0.519 | 0.600 | 1218 |
| Liriodendron | Tulip Tree | 0.546 | 0.486 | 0.514 | 1175 |
| Cladrastis | Yellowwood | 0.423 | 0.446 | 0.434 | 1110 |
| Sorbus | Mountain Ash | 0.551 | 0.458 | 0.500 | 1068 |
| Rhamnus | Buckthorn | 0.270 | 0.273 | 0.272 | 1065 |
| Cercidiphyllum | Katsura Tree | 0.726 | 0.701 | 0.713 | 1046 |
| Parrotia | Persian ironwood | 0.611 | 0.641 | 0.626 | 979 |
| Ailanthus | Tree of Heaven | 0.509 | 0.541 | 0.524 | 974 |
| Eriobotrya | Loquat | 0.527 | 0.578 | 0.551 | 963 |
| Morus | Mulberry | 0.345 | 0.447 | 0.390 | 816 |
| Styrax | Snowbell | 0.522 | 0.527 | 0.524 | 782 |
| Pseudotsuga | Douglass Fir | 0.601 | 0.538 | 0.567 | 766 |
| Maytenus | Mayten Tree | 0.570 | 0.669 | 0.615 | 759 |
| Metasequoia | Dawn Redwood | 0.667 | 0.532 | 0.592 | 631 |
| Cedrus | Cedar | 0.642 | 0.646 | 0.644 | 596 |
| Rhus | Sumac | 0.634 | 0.276 | 0.385 | 565 |
| Elaeagnus | Silverberry | 0.660 | 0.624 | 0.641 | 556 |
| Alnus | Alder | 0.416 | 0.321 | 0.362 | 527 |
| Salix | Willow | 0.487 | 0.267 | 0.345 | 509 |
| Sequoia | Coast Redwood | 0.581 | 0.686 | 0.629 | 474 |
| Juglans | Walnut | 0.209 | 0.408 | 0.277 | 463 |
| Myoporum | Myoporum | 0.170 | 0.237 | 0.198 | 456 |
| Casuarina | Sheoak | 0.889 | 0.832 | 0.860 | 394 |
| Chamaecyparis | False Cypress | 0.457 | 0.362 | 0.404 | 384 |
| Ostrya | Hop Hornbeam | 0.265 | 0.125 | 0.170 | 352 |
| Taxodium | Bald Cypress | 0.273 | 0.344 | 0.304 | 349 |
| Triadica | Chinese Tallow Tree | 0.570 | 0.738 | 0.643 | 347 |
| Citrus | Citrus | 0.415 | 0.420 | 0.418 | 333 |
| Corylus | Hazel | 0.420 | 0.222 | 0.291 | 333 |
| Abies | Fir | 0.554 | 0.205 | 0.299 | 273 |
| Nerium | Oleander | 0.130 | 0.485 | 0.205 | 260 |
| Albizia | Silk Tree | 0.478 | 0.473 | 0.476 | 258 |
| Larix | Larch | 0.663 | 0.567 | 0.611 | 233 |
| Phellodendron | Amur Cork Tree | 0.296 | 0.270 | 0.282 | 226 |
| Ilex | Holly | 0.262 | 0.363 | 0.305 | 223 |
| Persea | Avocado | 0.374 | 0.411 | 0.392 | 202 |
| Hibiscus | Hibiscus | 0.384 | 0.222 | 0.281 | 194 |
| Chionanthus | Fringe Tree | 0.183 | 0.104 | 0.132 | 183 |
| Taxus | Yew | 0.588 | 0.287 | 0.386 | 174 |
| Carya | Hickory | 0.097 | 0.045 | 0.062 | 132 |
| Tsuga | Hemlock | 0.310 | 0.247 | 0.275 | 89 |
| Castanea | Chestnut | 0.226 | 0.280 | 0.250 | 25 |

Table S4. Beta-regression models between weighted average F1 score and three diversity metrics (Shannon diversity, genus richness, and genus evenness) for tree genera across 23 study cities. Significant P-values (*P < 0.05*) are indicated in bold.

| Model: Weighted Average F1 Score ~ Shannon Diversity | Estimate | Std. Err | Z value | P |
| --- | --- | --- | --- | --- |
| Intercept | 2.0463 | 0.4796 | 4.267 | **<0.001** |
| Shannon Diversity | -0.4263 | 0.1466 | -2.908 | **0.004** |
| - | - | - | - | - |
| Model: Weighted Average F1 Score ~ Richness | Estimate | Std. Err | Z value | P |
| Intercept | 1.3391 | 0.3409 | 3.927 | **<0.001** |
| Richness | -0.0111 | 0.0055 | -2.008 | **0.045** |
| - | - | - | - | - |
| Model: Weighted Average F1 Score ~ Evenness | Estimate | Std. Err | Z value | P |
| Intercept | 3.1308 | 0.6992 | 4.478 | **<0.001** |
| Evenness | -3.1806 | 0.8974 | -3.544 | **<0.001** |

Table S5. ANOVA tables for classification performance across 17 genera by month, geography, and number of training images. Significant P-values (*p < 0.05*) are indicated in bold.

| Genus: *Acer* (Maple) | | | |
| --- | --- | --- | --- |
|  | df | F | P |
| Month | 11 | 3.164 | **0.001** |
| Region | 3 | 23.537 | **<0.001** |
| Training Image Count | 1 | 2.713 | 0.104 |
| Month * Region | 26 | 1.166 | 0.298 |
| Genus: *Fraxinus* (Ash) | | | |
|  | df | F | P |
| Month | 11 | 0.989 | 0.466 |
| Region | 3 | 7.763 | **0.002** |
| Training Image Count | 1 | 0.010 | 0.920 |
| Month * Region | 26 | 0.328 | 0.999 |
| Genus: *Quercus* (Oak) | | | |
|  | df | F | P |
| Month | 11 | 1.686 | 0.095 |
| Region | 3 | 9.546 | **<0.001** |
| Training Image Count | 1 | 2.407 | 0.125 |
| Month * Region | 27 | 0.994 | 0.489 |
| Genus: *Ulmus* (Elm) | | | |
|  | df | F | P |
| Month | 11 | 0.829 | 0.612 |
| Region | 3 | 6.866 | **<0.001** |
| Training Image Count | 1 | 7.109 | **0.001** |
| Month * Region | 25 | 0.736 | 0.798 |
| Genus: *Prunus* (Cherry/ Plum) | | | |
|  | df | F | P |
| Month | 11 | 1.692 | 0.094 |
| Region | 3 | 26.453 | **<0.001** |
| Training Image Count | 1 | 5.421 | **0.023** |
| Month * Region | 23 | 0.563 | 0.937 |
| Genus: *Tilia* (Basswood) | | | |
|  | df | F | P |
| Month | 11 | 1.855 | 0.070 |
| Region | 3 | 12.885 | **<0.001** |
| Training Image Count | 1 | 1.105 | 0.298 |
| Month * Region | 16 | 0.714 | 0.766 |
| Genus: *Pyrus* (Pear) | | | |
|  | df | F | P |
| Month | 11 | 0.703 | 0.729 |
| Region | 3 | 0.509 | 0.678 |
| Training Image Count | 1 | 1.254 | 0.269 |
| Month * Region | 20 | 0.896 | 0.594 |
| Genus: *Gleditsia* (Honeylocust) | | | |
|  | df | F | P |
| Month | 11 | 0.986 | 0.476 |
| Region | 3 | 0.761 | 0.523 |
| Training Image Count | 1 | 1.467 | 0.233 |
| Month * Region | 20 | 0.521 | 0.939 |
| Genus: *Malus* (Apples) | | | |
|  | df | F | P |
| Month | 11 | 0.828 | 0.614 |
| Region | 3 | 11.337 | **<0.001** |
| Training Image Count | 1 | 14.032 | **<0.001** |
| Month * Region | 14 | 0.945 | 0.523 |
| Genus: *Platanus* (Sycamore) | | | |
|  | df | F | P |
| Month | 11 | 0.365 | 0.962 |
| Region | 3 | 5.740 | **0.002** |
| Training Image Count | 1 | 0.335 | 0.561 |
| Month * Region | 19 | 0.743 | 0.754 |
| Genus: *Liquidambar* (Sweetgum) | | | |
|  | df | F | P |
| Month | 11 | 1.027 | 0.447 |
| Region | 2 | 3.159 | 0.056 |
| Training Image Count | 1 | 1.454 | 0.237 |
| Month * Region | 15 | 0.803 | 0.667 |
| Genus: *Pinus* (Pine) | | | |
|  | df | F | P |
| Month | 11 | 0.382 | 0.957 |
| Region | 3 | 0.715 | 0.548 |
| Training Image Count | 1 | 0.051 | 0.823 |
| Month * Region | 25 | 0.589 | 0.923 |
| Genus: *Magnolia* (Magnolia) | | | |
|  | df | F | P |
| Month | 11 | 0.430 | 0.930 |
| Region | 3 | 8.094 | **<0.001** |
| Training Image Count | 1 | 3.983 | 0.055 |
| Month * Region | 13 | 0.956 | 0.513 |
| Genus: *Picea* (Spruce) | | | |
|  | df | F | P |
| Month | 8 | 0.402 | 0.910 |
| Region | 2 | 1.037 | 0.367 |
| Training Image Count | 1 | 2.708 | 0.110 |
| Month * Region | 10 | 0.724 | 0.696 |
| Genus: *Ginkgo* (Ginkgo) | | | |
|  | df | F | P |
| Month | 11 | 0.653 | 0.771 |
| Region | 3 | 2.741 | 0.058 |
| Training Image Count | 1 | 4.907 | **0.034** |
| Month * Region | 14 | 0.405 | 0.963 |
| Genus: *Celtis* (Hackberry) | | | |
|  | df | F | P |
| Month | 11 | 0.962 | 0.502 |
| Region | 3 | 1.786 | 0.174 |
| Training Image Count | 1 | 9.627 | **0.005** |
| Month * Region | 13 | 0.657 | 0.784 |
| Genus: *Crataegus* (Hawthorn) | | | |
|  | df | F | P |
| Month | 11 | 0.890 | 0.563 |
| Region | 3 | 6.365 | **0.003** |
| Training Image Count | 1 | 0.384 | 0.542 |
| Month * Region | 19 | 0.764 | 0.722 |

Table S6. Percent of inventory trees matched with at least one Google Street View image within the 20-m search radius by genus.

| **Genus** | **Percent tree matched** |
| --- | --- |
| casuarina | 93.1% |
| cinnamomum | 92.0% |
| melaleuca | 87.2% |
| cupaniopsis | 86.8% |
| platanus | 85.9% |
| pistacia | 84.7% |
| gleditsia | 83.7% |
| jacaranda | 83.7% |
| liquidambar | 81.4% |
| eucalyptus | 81.1% |
| schinus | 80.4% |
| acer | 79.3% |
| ulmus | 78.8% |
| tilia | 78.0% |
| zelkova | 76.9% |
| ficus | 76.1% |
| magnolia | 75.9% |
| ailanthus | 75.3% |
| pyrus | 75.1% |
| metrosideros | 74.0% |
| fraxinus | 73.4% |
| quercus | 71.3% |
| sequoia | 70.1% |
| ceratonia | 68.8% |
| arbutus | 68.3% |
| styphnolobium | 68.1% |
| fagus | 68.0% |
| triadica | 68.0% |
| catalpa | 66.3% |
| cedrus | 66.2% |
| cercidiphyllum | 64.7% |
| lagerstroemia | 64.6% |
| celtis | 64.0% |
| podocarpus | 63.9% |
| callistemon | 63.1% |
| olea | 61.2% |
| lophostemon | 61.0% |
| pinus | 61.0% |
| tristaniopsis | 60.4% |
| koelreuteria | 59.0% |
| pseudotsuga | 59.0% |
| maytenus | 58.8% |
| syagrus | 58.4% |
| populus | 58.2% |
| ginkgo | 57.9% |
| liriodendron | 56.1% |
| albizia | 55.8% |
| picea | 55.8% |
| aesculus | 54.9% |
| prunus | 54.6% |
| carpinus | 54.1% |
| ligustrum | 51.7% |
| gymnocladus | 51.4% |
| washingtonia | 51.4% |
| pittosporum | 49.8% |
| eriobotrya | 48.7% |
| betula | 48.3% |
| acacia | 46.4% |
| robinia | 45.6% |
| malus | 45.0% |
| alnus | 43.8% |
| nyssa | 42.9% |
| phoenix | 42.8% |
| cercis | 42.4% |
| styrax | 39.4% |
| crataegus | 38.3% |
| parrotia | 38.0% |
| metasequoia | 37.6% |
| myoporum | 37.3% |
| cornus | 37.3% |
| sorbus | 37.3% |
| abies | 36.7% |
| elaeagnus | 36.1% |
| cupressus | 33.9% |
| amelanchier | 32.5% |
| cladrastis | 32.1% |
| morus | 31.4% |
| taxodium | 31.0% |
| tsuga | 30.8% |
| syringa | 30.1% |
| juglans | 29.8% |
| nerium | 29.7% |
| ilex | 28.6% |
| juniperus | 27.1% |
| corylus | 26.4% |
| thuja | 25.8% |
| larix | 24.7% |
| rhus | 21.2% |
| phellodendron | 20.5% |
| salix | 19.8% |
| persea | 18.8% |
| chamaecyparis | 17.2% |
| carya | 16.7% |
| citrus | 13.6% |
| hibiscus | 10.0% |
| rhamnus | 8.8% |
| chionanthus | 5.9% |
| ostrya | 5.7% |
| taxus | 4.5% |
| castanea | 0.0% |
